# Frontal cortex encodes action goals and social context in freely moving and socially interacting macaques

**DOI:** 10.1101/2025.10.21.683128

**Authors:** Jacopo Baldi, Eloise Disarbois, Andres C. Mendez, Giorgio Cappellaro, Giulia Annicchiarico, Marco Bimbi, Gino Coudé, Jean-René Duhamel, Pier Francesco Ferrari

**Affiliations:** Institut des Sciences Cognitives Marc Jeannerod, CNRS, 67 Blvd. Pinel, Bron, France; Université Lyon 1 Claude Bernard, Lyon, France; Dipartimento di Medicina e Chirurgia, Università di Parma, Via Volturno 39, 43125, Parma, Italy

## Abstract

Classic studies of goal-directed behavior in primates have investigated frontal cortical circuits under restraint conditions, leaving their role in natural social contexts largely unknown. We wirelessly recorded neural activity from freely moving macaques to study how frontal areas encode actions during naturalistic interactions. Neurons in ventral premotor and ventrolateral prefrontal cortex distinguished identical motor acts, such as grasping, depending on whether they occurred during foraging or grooming, indicating sensitivity to social meaning. Some units were selectively tuned to socially directed goals, while population analyses showed both regions flexibly integrated motor plans with social context. These results provide direct evidence that these regions, long considered key motor hubs, also encode the social dimension of action, redefining current models of frontal lobe function.

**One-Sentence Summary:** Freely-moving, social-interacting macaques reveal new properties of frontal circuits in encoding motor goals and social context.

## Main Text

For decades, systems neuroscience has relied on non-human primates to investigate the neural bases of goal-directed actions, exploiting their strong anatomical and functional homologies with humans. Classic neurophysiological work with macaques under highly controlled conditions has been instrumental in defining the properties and computations of neurons within specific regions of the frontal lobe. In particular, studies of the ventral premotor cortex (PMv; area F5) have revealed neurons that integrate visual, somatosensory, and motor information to encode specific goals such as grasping, which in some cases, occurs regardless of the effector used (i.e. the hand or the mouth) or even during tool use (*1–4*). This region receives convergent input from the anterior intraparietal area, providing visual information about object properties and affordances (*5–7*), enabling sensorimotor transformations for action (*8*). Beyond object features, some F5 visuomotor neurons are sensitive to others’ actions, intentions, and social goals (*9–15*), thus showing a rich modulation of F5 neurons relative to the context.

The ventrolateral prefrontal cortex (vlPFC; areas 46v, 12r, 45a) complements F5 by supporting higher-order aspects of goal-directed behavior. It contributes to action planning, motor sequencing; (*16–18*), rule-based behavior (*19*, *20*), and visuomotor coordination (*21*, *22*). It also processes complex social stimuli including audiovisual integration (*23*) and the perception of social interactions (*24*). Importantly, vlPFC is strongly interconnected with F5 (*25*, *26*), positioning it as a strategic hub for integrating motor and cognitive components of action control.

These discoveries have profoundly shaped our view of frontal lobe functions. However, much of this knowledge stems from experiments conducted under highly constrained laboratory conditions (i.e. using head-restrain and primate chairs), where monkeys are tested in standardized tasks, requiring stereotyped arm/hand actions. While powerful, these paradigms provide only a partial view, leaving unexplored how the frontal lobe encodes motor behaviors in the rich contexts of natural actions and social interactions. Recent advances in neuroscience have opened new avenues for studying brain function under ecologically valid conditions, thanks to wireless recording systems in freely moving animals. Though still-emerging, this approach is already yielding important results. Recent methodological and conceptual neuroscientific advances are now challenging the assumption that neural mechanisms observed under restrained conditions can fully generalize to real-world behavior. The use of wireless recording systems in freely moving animals now offers a powerful alternative, enabling the study of brain activity in the very conditions for which it evolved: ecological, dynamic, and interactive. For example, wireless recordings in flying bats have revealed hippocampal dynamics that fundamentally reshape our understanding of how neural circuits support behavior in natural environments (*27*).

The technological leap required to record neural activity in freely moving macaques has been particularly demanding, but it holds exceptional promise for expanding our understanding of brain functions, especially in terms of translating animal models to human cognition and behavior. Very recently, wireless studies in macaques have shown that premotor and parietal circuits encode distant goals rather than mere motor acts (*28*), and that neuronal activity in F5 diverges between identical actions performed under restrained versus unrestrained conditions, with neural patterns in the naturalistic context generalizing to the laboratory but not vice versa (*29*). Crucially, some units also show mixed selectivity for a wider variety of behaviors in the free-moving condition, suggesting that ecological paradigms can uncover previously hidden dimensions of neural coding. Recordings during natural social interactions likewise challenge established views of functional specialization in prefrontal and temporal areas (*30*). However, despite these pioneering efforts, the study of the motor system in real social situations remains significantly absent, thus preventing a full comprehension of how neural circuits support action in ecologically meaningful, socially interactive contexts.

Here we build on this emerging perspective to investigate how frontal regions, specifically PMv and vlPFC, represent motor goals in freely moving and socially interacting macaques. By employing wireless neural recordings in naturalistic settings, we examine how social and non-social contexts modulate goal-directed actions, and how neurons in the frontal cortex flexibly integrate environmental and social information. In doing so, we aim to provide an ecologically grounded account of frontal lobe function, bridging the gap between controlled experimental findings and the complexity of natural behavior. Given the anatomical and functional links between regions encoding social signals and those involved in action goals, we hypothesized that neural activity in the frontal lobe would differ when similar behaviors are performed in social versus non-social contexts. To test this, we focused on two distinct natural behaviors: 1) foraging, which is primarily individual and object/food-related, and 2) grooming, which is inherently social. By recording neural activity from both PMv and vlPFC, we can directly compare how these regions encode motor goals under different social conditions. We predicted that PMv would provide more specific representations of motor goals, given its proximity to motor output, whereas vlPFC would more strongly integrate social dimensions through its higher-order contextual functions.

### Subjects and Setting

We investigated the brain activity of two female monkeys (M1 and M2, *Macaca mulatta*) while they were free to move and interact with their respective female partners (non-implanted) and objects and food present in the environment of the EthoControl Room (ECR) (Fig. 1A, Supp. Mat., Fig S1E). In the respective pairs, M1 was the submissive while M2 was the dominant monkey (see Methods for how we defined the hierarchy). The ECR is a platform for wireless electrophysiological and behavioral recording in non-human primates, consisting of a room enclosed by anti-reflective glass panels, and surrounded by fixed and motorized cameras. The fixed cameras provided real-time tracking of the subject’s position in space, and the motorized cameras used that information to reorient and follow the monkey for high-resolution behavioral views. The camera system was controlled by EthoLoop software (*31*), while DeuteronTech software managed the Neural Logger device (Supp. Mat., Fig S1D); inter-software communication ensured precise synchronization of behavioral and neural data. We recorded neuronal activity from sets of 128 electrodes chronically implanted in the pre- and post-arcuate sulcus of two rhesus monkeys (Fig. S1A, B). We simultaneously recorded extracellular activity via four 32-channel floating multielectrode arrays (FMAs, Supp. Mat., Fig. S1G, H), using two arrays implanted in ventrolateral prefrontal cortex (vlPFC Brodmann areas 46v and 45a) and two in ventral premotor cortex (PMv, specifically area F5), and a custom wireless headstage to interface the Omnetics connectors of the chronic implant with the Neurologger. A 3D-printed plastic box (Supp. Mat., Fig. S1F) was installed and secured on the monkey’s head to protect the device during free-moving sessions. Data were collected across three sessions per monkey, chosen to capture a wide variety of naturally occurring behaviors.

**Figure 1:**
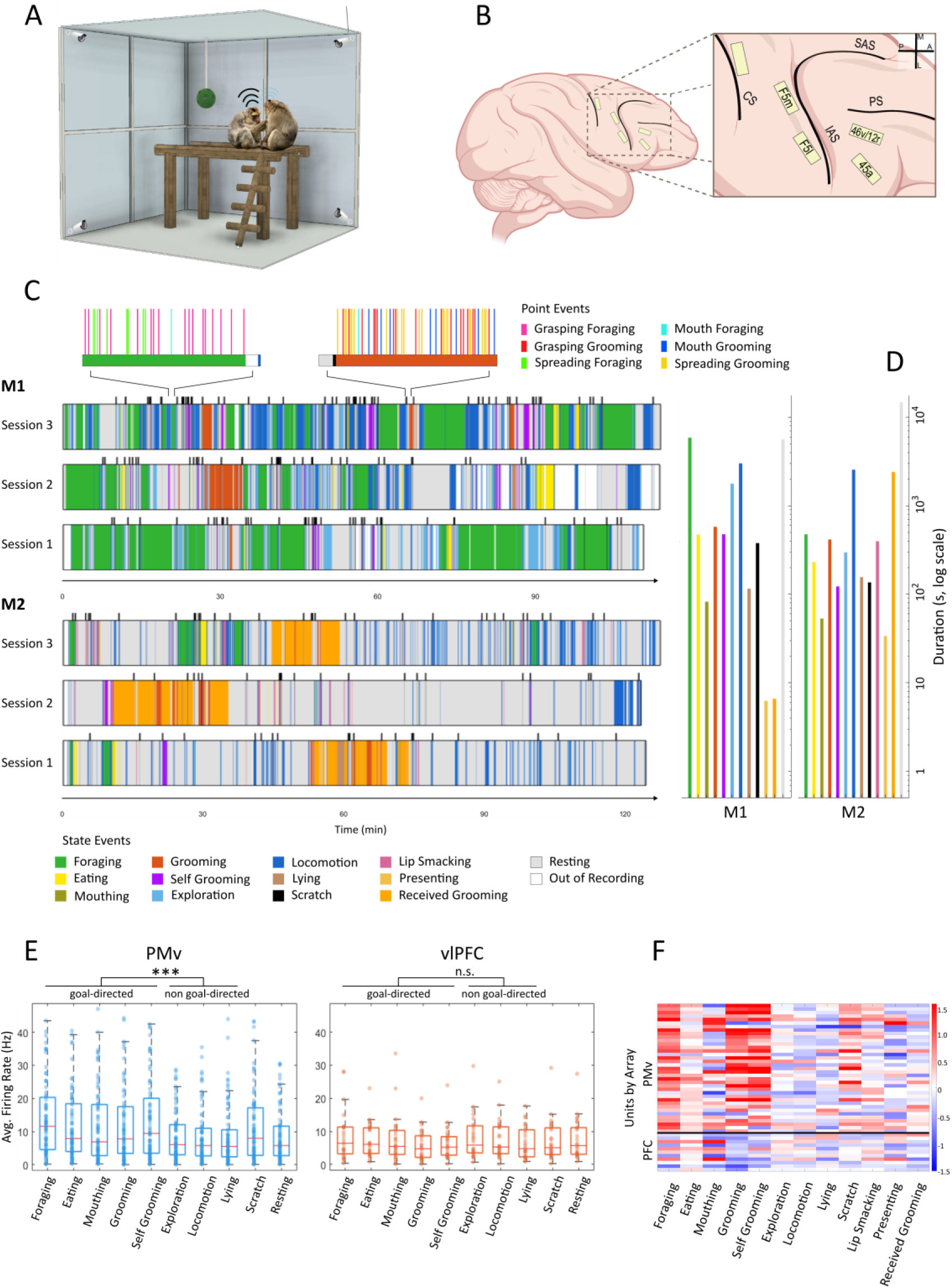
Experimental setting, implant site, behavioral analysis and state-based analysis. (A) Schematic illustration of the ECR showing a monkey equipped with the wireless recording device. (B) Zoomed schematic of the frontal lobe implant site indicating the placement of PMv and vlPFC arrays (created with BioRender.com). (C) Session-wide ethograms for each recording session showing the sequence of behavioral states. M1 above and M2 below. Upper section: zoomed-in section of a single session illustrating point events during foraging (left) and grooming (right). (D) Quantification of total time spent in each behavioral state (time budget) on logarithmic scale. (E) Boxplot visualization of firing rates of units pooled across monkeys during different behavioral states, and statistical comparison between goal-directed and non-goal-directed groups of behaviors. (F) Cohen’s D effect size metric of units’ firing rate distributions during each behavioral class against resting state. Blue means lower firing rate compared to rest, red means higher.

Spike sorting was performed first separately for each session to establish a baseline of how many units could be identified per session (Table 1: Total neurons across sessions: M1; N=206; M2; N=155), and then on the concatenated recordings to analyze their modulation across behaviors pooled from all recording sessions. Stable single- and multi-unit isolation throughout the concatenated recording was confirmed based on waveform consistency and firing-rate stability, selecting only units that could be reliably tracked. The loss in unit yield after concatenation was relatively small (Table 1: Concatenated): for monkey M1, from an overall pool of 206 identified units, the number of units decreased from an average of 69 per session to 57 (≈17% reduction), while for monkey M2, from a total of 155, the number of units decreased from an average of 52 to 48 (≈8% reduction). For both monkeys, the number of units in the arrays in PMv cortex far surpassed that in vlPFC, with M1 having 42 putative units in PMv versus 15 in vlPFC and M2 having 37 in PMv versus 11 in vlPFC (Table 1).

**Table 1:** Number of putative units detected with spike sorting during each session (S1-3 of each monkey), total sum of the three, and concatenated, per each monkey (M1 and M2) and recording area (PMv and vlPFC).

|  |  | S1 | S2 | S3 | Total | Avg. | Concatenated |
| --- | --- | --- | --- | --- | --- | --- | --- |
| M1 | PMv | 51 | 46 | 40 | 137 | 46 | 42 |
|  | vIPFC | 21 | 25 | 23 | 69 | 23 | 15 |
|  | Total | 72 | 71 | 63 | 206 | 69 | 57 |
| M2 | PMv | 38 | 39 | 41 | 118 | 39 | 37 |
|  | vIPFC | 11 | 14 | 12 | 37 | 12 | 11 |
|  | Total | 49 | 53 | 53 | 155 | 52 | 48 |

### Free-moving monkeys display diverse, naturalistic behaviors under wireless recording conditions

During the recording sessions each monkey and her partner exhibited a wide range of social and non-social natural behaviors, predominantly exploration of the cage, foraging, and grooming (Fig. 1C, Supp. Mat., Fig. S1C). In order to minimize experimenter influence on the natural behavior, we decided to not intervene in any way to encourage certain behaviors, except for scattering different pieces of food around the cage to allow foraging, leaving the monkeys complete freedom to engage in any activity and interact with each other. The behavioral repertoire that both pairs of monkeys expressed was qualitatively the same as observed in their home enclosure, indicating no detectable impact of the recording apparatus on their behavior. See Methods for a detailed breakdown of how we defined each behavior in this space. Two experimenters trained on macaque behaviors coded both state-based behaviors – such as foraging or grooming – and point events – single frames representing a point in an action sequence, such as the moment the fingers of the monkey touch a piece of food during a reaching-and-grasping sequence. The drawback of this approach, as can be seen in Figure 1D, is a heavily skewed distribution of the behaviors they expressed between different sessions, with some classes of behaviors being absent during some sessions and overrepresented during others (Supp. Mat., Table S1 reports exact numbers of time spent in each behavior). To ensure every behavior class was available for neural analyses, we concatenated all three sessions per monkey, treating them as a single continuous dataset. There were substantial qualitative differences in the overall level of activity and behaviors expressed by the two monkeys. M1 is younger and more active than M2, and spent substantial portions of time resting and foraging for food (30.6% Resting and 31.9% Foraging of the entire sessions, respectively; Suppl. Mat., Table S1), a smaller but significant portion exploring the cage (16.4% Locomotion), and much smaller periods grooming the partner or herself (3.1% Grooming, 2.6% Self-grooming). Due to a strongly skewed dominance relationship, she performed grooming with no reciprocation at all from her partner. In a very different pattern, M2 preferred to spend the vast majority of her time resting on the platform in the cage (66.9% Resting) and dedicated a much smaller fraction (2.2%) to foraging for food. She spent a similar amount of time as M1 exploring the cage (11.6% Locomotion) and grooming her partner (1.9% Grooming), but unlike M1, being the dominant, she received a considerable amount of grooming from her subordinate (11.0% Received grooming). She did very little self-grooming throughout the sessions (0.6% Self-grooming). In line with these differences, presenting was done more by M2 (0.15% vs 0.03% in M1), whereas exploratory behaviors were much more frequent for M1 (9.6% Exploration vs 1.4% in M2), while lipsmacking was exclusively shown by M2 (1.8%). Smaller behaviors in terms of percentage, like Mouthing (0.45% vs 0.24%), Lying (0.63% vs 0.71%), Scratching (2.05% vs 0.62%), and Eating (2.6% vs 1.1%), did not show clear differences across monkeys.

Each session was different from the others, with a very unstructured sequencing of periods of resting, exploration, foraging and social interactions with the partner. One of the regularities that seems to emerge is that in the case of M2 most of the social interactions (giving and receiving grooming) were concentrated in a single isolated long bout in each session. Figure 1C (upper section) shows the sequence of point events that happen during two sample snippets of complex behavior, one during foraging (upper-left section) which shows a sequence of spreading in search for food and grasping to bring it to the mouth, and the other during grooming (upper-right section) which shows the same pattern replicated on the fur: spreading with the hands, followed by grasping the target with either a hand or the mouth. Each bout of foraging or grooming state contained many instances of single point-events (Supp. Mat., Table S2 reports exact frequency of events). A key difference between M1 and M2 was that M2 never used her mouth to grab food during foraging or particles during grooming, entirely preventing us from analyzing that specific behavior in this monkey. We also removed from the analysis behaviors that were produced for a total of less than 10s across the three sessions of each monkey (i.e. Presenting and Received Grooming for M1), because the sample was not enough for statistical testing.

### State-based population analysis shows different levels of activity for goal-directed classes of behaviors

The first axis of analysis aimed to assess whether different behavioral states were associated with macroscopic differences in neuronal activity levels, measured as the average firing rate during each behavior. Figure 1E shows the average firing rates of individual units (one dot per column) across behavioral classes. To increase statistical power and capture generalizable effects, we pooled the neuronal populations of both monkeys and compared firing-rate distributions across behavioral categories. In the PMv area (Fig. 1E, left panel), different behaviors exhibited distinct population activity levels. Given the known role of the premotor cortex in goal-directed motor acts, we expected PMv neurons to display higher activity during goal-directed behaviors.

To test this, we grouped behavioral states into goal-directed (Foraging, Eating, Mouthing, Grooming, and Self-Grooming) and non–goal-directed (Exploration, Locomotion, and Lying) categories (*32*) (see Methods section for the distinction), and compared their firing-rate distributions using a nonparametric permutation test (100,000 iterations). When pooling data across both monkeys, PMv activity was significantly higher during goal-directed behaviors compared to non–goal-directed ones (p = 5 × 10⁻⁵), while vlPFC showed no significant difference between the two groups (p = 0.749, Fig. 1E, right panel). Similar results were obtained when the two monkeys were analyzed separately (Supp. Mat., Fig. S2B, C). These findings indicate that premotor—but not prefrontal—activity preferentially reflects goal-directed behaviors, consistently across animals.

To directly compare our dataset with the findings of Testard et al. (*30*), who investigated neural modulation in the vlPFC of freely moving monkeys using Cohen’s D, we adopted the same metric to quantify the strength and direction of activity changes between behaviors and resting state. This parallel approach allowed us to test the reproducibility of their results in an independent dataset recorded under highly similar ethological conditions.

For each monkey analyzed separately, Figure 1F (results of M2) displays the Cohen’s D coefficients for all recorded units and behavioral states (Supp. Mat., Fig. S2A shows results for M1). Positive values (red) indicate higher firing rates during the behavior compared to rest, while negative values (blue) indicate suppression. Units are ordered vertically by cortical array, with premotor arrays shown above and prefrontal arrays below, separated by a black horizontal line.

In both animals, we observed strong positive modulation in premotor cortex (PMv) during goal-directed behaviors such as Foraging, Eating, and Mouthing (see Methods for the definition of each behavior), consistent with the region’s involvement in motor control of hand and mouth actions. Other not goal-directed behaviors involving the movements of the arm and the hand (e.g. locomotion and exploration) showed a less robust discharge than goal-directed ones. Conversely, vlPFC units showed predominantly negative modulations during Grooming and Self-Grooming, in line with the pattern of activity reported by Testard et al. (*30*). Similar area-specific effects were also evident for Scratching (positive in PMv, negative in vlPFC) and Received Grooming (mostly negative in vlPFC). Other behavioral states elicited weaker or mixed effects. These results confirm that the modulation profiles originally described by Testard and colleagues can be reproduced in separate set of freely interacting subjects, extending the generality of those findings across individuals.

### Context-dependent neural tuning in PMv and vlPFC

To examine how neural activity distinguishes social and non-social goal-directed behaviors, we analyzed single-unit activity in PMv and vlPFC during naturally occurring foraging (non-social) and grooming (social) episodes. Spike-sorted units from concatenated sessions were aligned to behavioral events within ±500 ms windows, and spikes were binned at 1 ms resolution for subsequent analyses. Figure 2 shows raster plots and peristimulus time histograms (PSTHs) of units recorded in M1 (Supp. Mat., Fig. S3 for example units from M2) for six ethologically matched behaviors—grasping, spreading, and using the mouth to grab something—performed in both foraging and grooming contexts.

**Figure 2.**
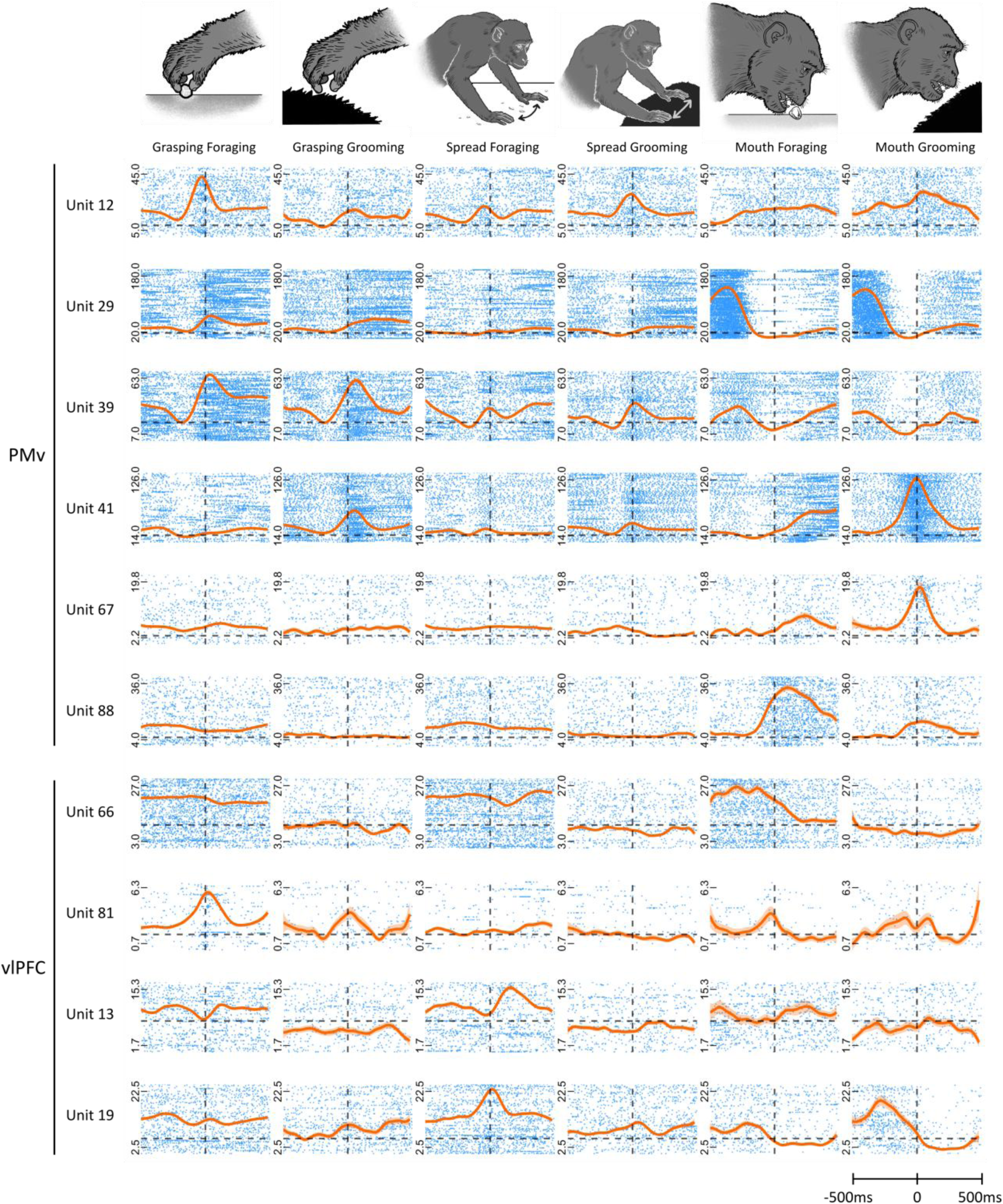
Single-unit activity from M1 aligned to point events. Raster plots and overlaid peristimulus time histograms (PSTHs) for multiple units. Rows correspond to individual units (names and area on the left); columns correspond to behaviors (icon and name at the top). Spike trains are taken in a window of 1s centered on the behavioral events as described in Methods.

Units in PMv exhibited clear tuning to specific motor goals. Many showed robust, time-locked responses similarly during both social actions and their non-social counterparts (e.g., Units 12, 29, 39, 81). Notably, some PMv units (e.g., Unit 41) responded more strongly to grooming-related behaviors than to their foraging counterparts, despite highly similar motor execution, indicating modulation by the social context. Other units (e.g., Unit 12) fired more strongly for grasping food than for grasping during grooming. Unit 39 fired for any type of hand grasping independently from the context. We also found mouth specific neuronal activity depending on the context (e.g., Units 67 and 88). Specific inhibited responses, while not common, were found as depicted in the example of Unit 29, where the neuron showed robust inhibited response during grasping with the mouth independently from the context. In the case of unit 13 instead, the inhibition was time-locked only for grasping during foraging, in contrast to an excited response for spreading during foraging, and an overall lower stable firing rate for the grooming behaviors. In contrast, some units recorded from vlPFC (e.g., Unit 19) showed weaker and more diffused modulations across all conditions. Unit 66 is an example of units that did not have a strong time-locked response to the events but had a big change in baseline activity during the relative states (e.g., much higher stable firing rates during foraging behaviors than grooming). This difference suggests that, in the control of natural behaviors, PMv may play a more prominent role in transforming contextual and sensory information into sensorimotor representations that can be implemented and relayed to the primary motor cortex and downstream corticospinal pathways. In contrast, vlPFC may contribute more variably across individuals, possibly reflecting its involvement in higher-order aspects of behavior such as attentional modulation, decision-making, or social context processing.

### Entropy-based approach uncovers extensive multiplexing of neural activity

Next, we aimed to identify, for each behavior, which units showed event-related modulation and to quantify each unit’s degree of selectivity and multiplexing. A key challenge when using free-moving recordings is establishing a reliable baseline. Most standard approaches to neural data analysis (e.g., ANOVA, GLMs) rely on comparing activity in a condition of interest to a clearly defined baseline, typically the pre-stimulus period in trial-based designs. We initially defined a unit’s baseline firing rate as its mean spike count over the entire session. To validate this choice, we also computed each unit’s mean firing rate during periods of monkey inactivity; these two measures were nearly identical across all units, so we adopted the session-wide average as the baseline. However, firing rates fluctuated substantially over time—even within comparable behavioral states—making any localized “control” period too variable to be reliable.

To circumvent this, we developed an information-theoretic approach based on Shannon entropy: if a unit’s spike pattern around an event carried significant information, we considered it task-related. Specifically, we computed the entropy of each unit’s real spike trains in the ±500 ms event window and compared it to the distribution obtained from spike trains with timestamps randomly shuffled within the same windows. Using the Wilcoxon signed-rank test (α = 0.05), units were classified as task-related if their observed entropy was significantly lower than that of the shuffled data. In order to validate this approach, we tested on a separate dataset obtained under restrained conditions (which provided a valid baseline) the degree of agreement between the new entropy-based method and a standard Generalized Linear Model (GLM) to measure differences between condition and baseline (Supp. Mat., Fig. S5). For each unit, and for each behavior class (hand grasping, hand spreading, and mouth grasping), we determined whether its modulation was significant in the foraging context, grooming context, both, or neither. Both animals showed a high prevalence of multi-behavior modulation in PMv (M1: 88.1%; M2: 91.9% for ≥3 behaviors). vlPFC showed inter-animal variability: M1 vlPFC units were strongly multi-behavior tuned (80.0% ≥3 behaviors; mean = 4.33 behaviors/unit), whereas M2 vlPFC included many unmodulated units (54.5% with 0 behaviors; mean = 1.55 behaviors/unit). Premotor neurons tended to be broadly tuned: ≈90% of PMv units were modulated for three or more behaviors, compared with ≈65% of vlPFC units (combined across M1 and M2) (Supp. Mat., Table S3). We then examined the selectivity of PMv and vlPFC units for foraging and grooming contexts (Fig. 3). Units were classified as modulated only during foraging (blue), only during grooming (red), during both behaviors (purple), or during neither (green). Across both monkeys, most PMv units were modulated for foraging behaviors or both (mean across PMv pies ≈ 92.0%), whereas vlPFC showed differences between subjects. In M1, all three vlPFC pies showed 86.7%, whereas M2 vlPFC pies were much lower (45.5% and 36.4%, mean ≈ 40.9%). Given the small absolute number of vlPFC units, it is difficult to draw meaningful conclusions about regions differences.

**Figure 3.**
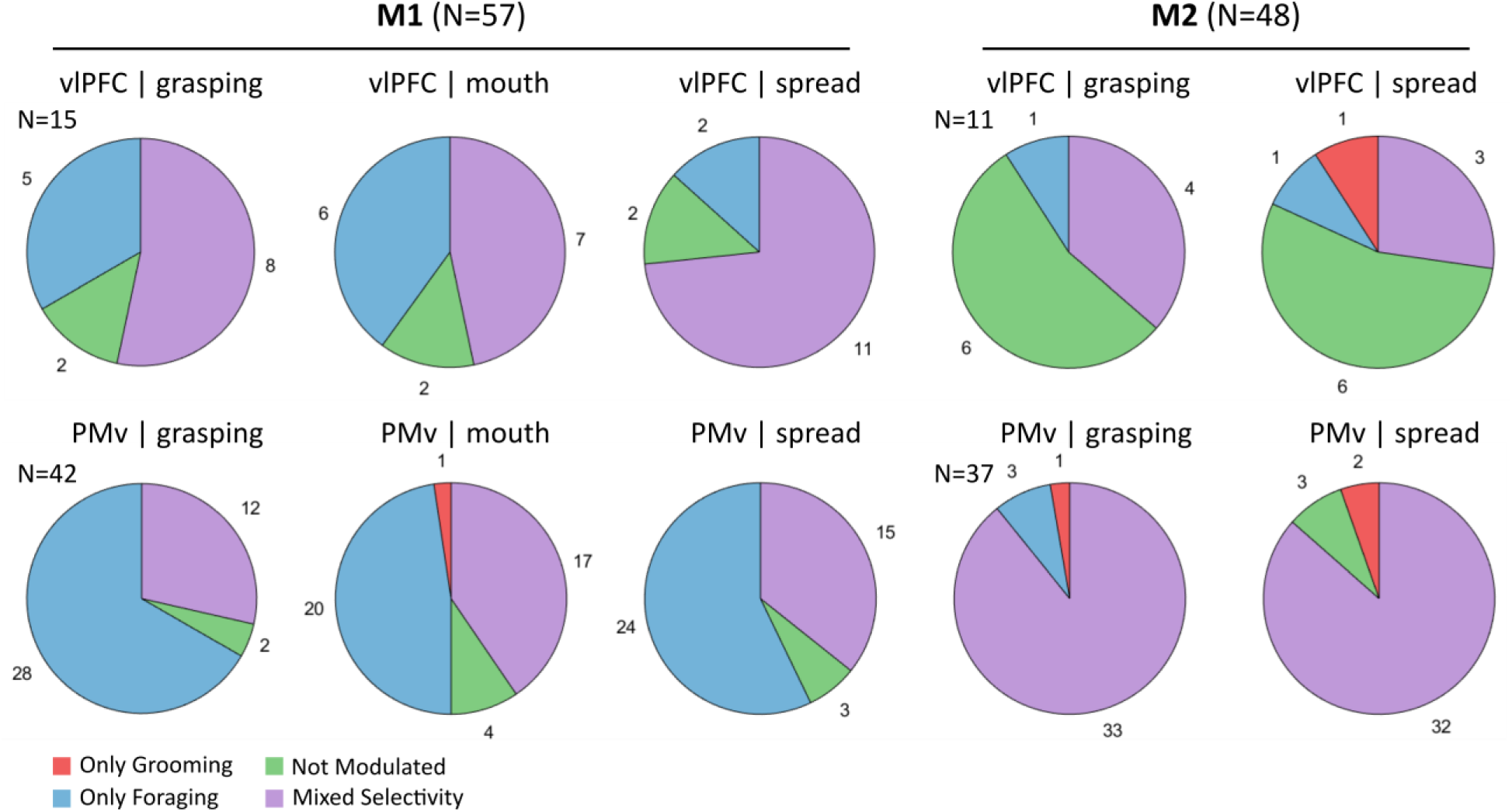
Proportion of units modulated by behaviors. Pie charts showing the numbers of units significantly modulated during each paired behavioral class. Colors indicate modulation type: blue = foraging only, red = grooming only, purple = both, green = unmodulated. Separate charts are shown for vlPFC and PMv (top row and bottom row, respectively), and for each monkey (M1 on the left, M2 on the right).

### Direction of modulation reveals stable excitatory tuning in PMv

We next analyzed the direction of response for units modulated by each behavior (Table 2). For each unit, we quantified the area between its PSTH response and the baseline firing rate, and determined whether the modulation was predominantly positive, negative, or biphasic (i.e., containing both positive and negative components). We used the biphasic class because the responses of some units during specific behaviors (i.e. Unit 19 in Fig. 2 for Mouth grooming having an upward modulation just before the action and a downward one just after it) could not be easily characterized as being just positive or negative, but both directions of modulation are plausibly relevant for its role in the network. In M1 PMv, foraging-related hand grasping was almost exclusively positively modulated (36 out of 40 units), with few mixed (n=3) and only one negative response. Similar patterns were observed for hand spreading and mouth grasping, with positive modulation dominating and mixed responses present in a minority of units. Grooming-related modulation in M1 PMv was more variable, with mixed responses emerging across behaviors (e.g., 4 out of 11 hand spreading units). M1 vlPFC units showed fewer mixed responses overall, with foraging-related hand grasping and spreading primarily positive, while grooming elicited a small number of negative or mixed responses. In M2, PMv units were again largely positively modulated for both foraging and grooming behaviors, with only occasional mixed responses for hand spreading and mouth grasping. M2 vlPFC units were sparse, with small numbers distributed across positive, negative, and mixed categories. Across both animals and cortical areas, PMv units predominantly exhibited positive modulation, whereas grooming-related responses and vlPFC units showed a more heterogeneous mix, suggesting more complex or context-dependent encoding in those conditions.

**Table 2:**
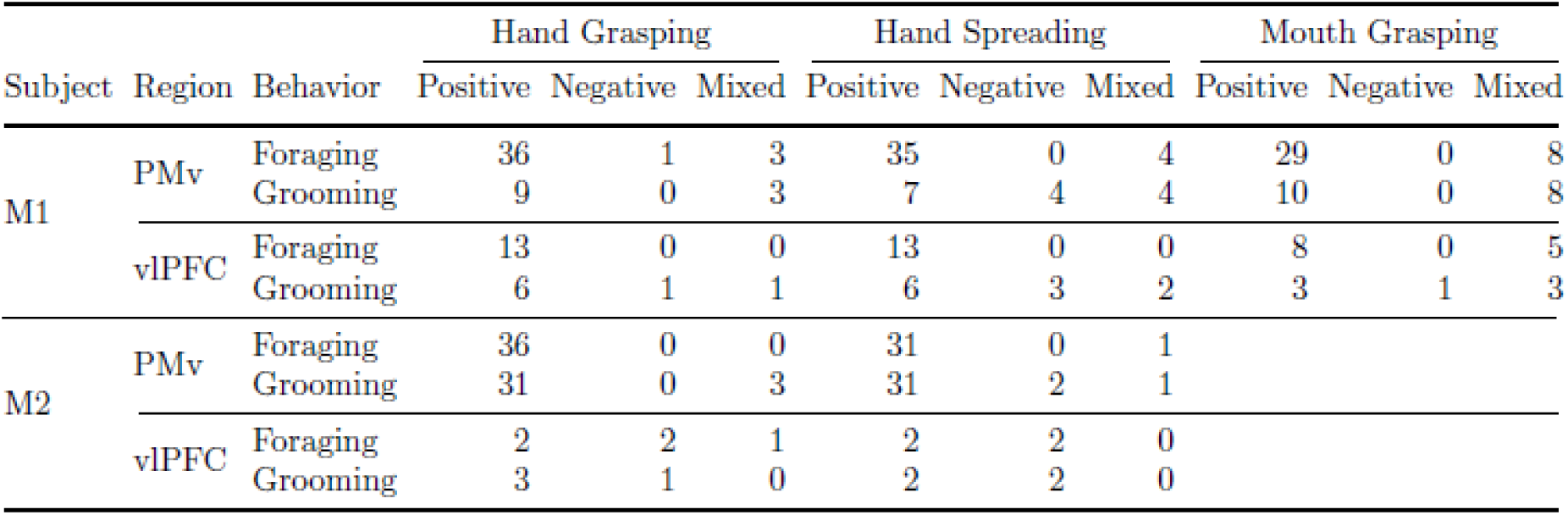
Directionality of response for units modulated by different behaviors in premotor (PMv) and prefrontal (vlPFC) cortices of the two subjects (M1 and M2) during foraging and grooming behaviors. Responses are categorized as positive, negative, or mixed for each of the three distinct action types (Hand Grasping. Hand Spreading, Mouth Grasping), and subdivided by the context of the response (Foraging vs Grooming). The response of M2 for Mouth Grasping behaviors is absent because the monkey does not produce that behavior.

### Context discrimination emerges strongly and foraging preference dominates the neural encoding

Next, we investigated whether units modulated for the same behavioral class during both social (grooming) and non-social (foraging) conditions (purple sections in Fig. 4) could discriminate between contexts, using the cluster-based permutation test (CBPT, see Methods for implementation). This method allowed direct comparison of the average response of a unit between conditions (Table 3). In M1, most units with mixed selectivity in both vlPFC and PMv were discriminating (vlPFC: 50–100%; PMv: 92–100%). Likewise, in PMv of M2 most units discriminated (94% for grasping, 69% for spreading), whereas vlPFC showed lower proportions (50% and 33%). Overall, units jointly modulated by both behaviors formed the largest proportion and are likely informative for distinguishing social vs. non-social contexts. The final classification layer assessed context preference in units that were selective for both states. Using the same area-based measure described above, we compared the absolute magnitude of modulation between foraging and grooming responses. When the difference between responses was less than 10%, we decided to classify the units as having no preference (though they could still be discriminating per CBPT), a conservative threshold to avoid misinterpreting minor fluctuations. In both M1 and M2, over 50% of PMv units showed a bias toward foraging across all behaviors, with the remainder split between grooming preference, equal responses, or non-discriminating. vlPFC units did not show a clear pattern of preference. Overall, PMv units exhibited the strongest and most consistent context preference for foraging.

**Figure 4.**
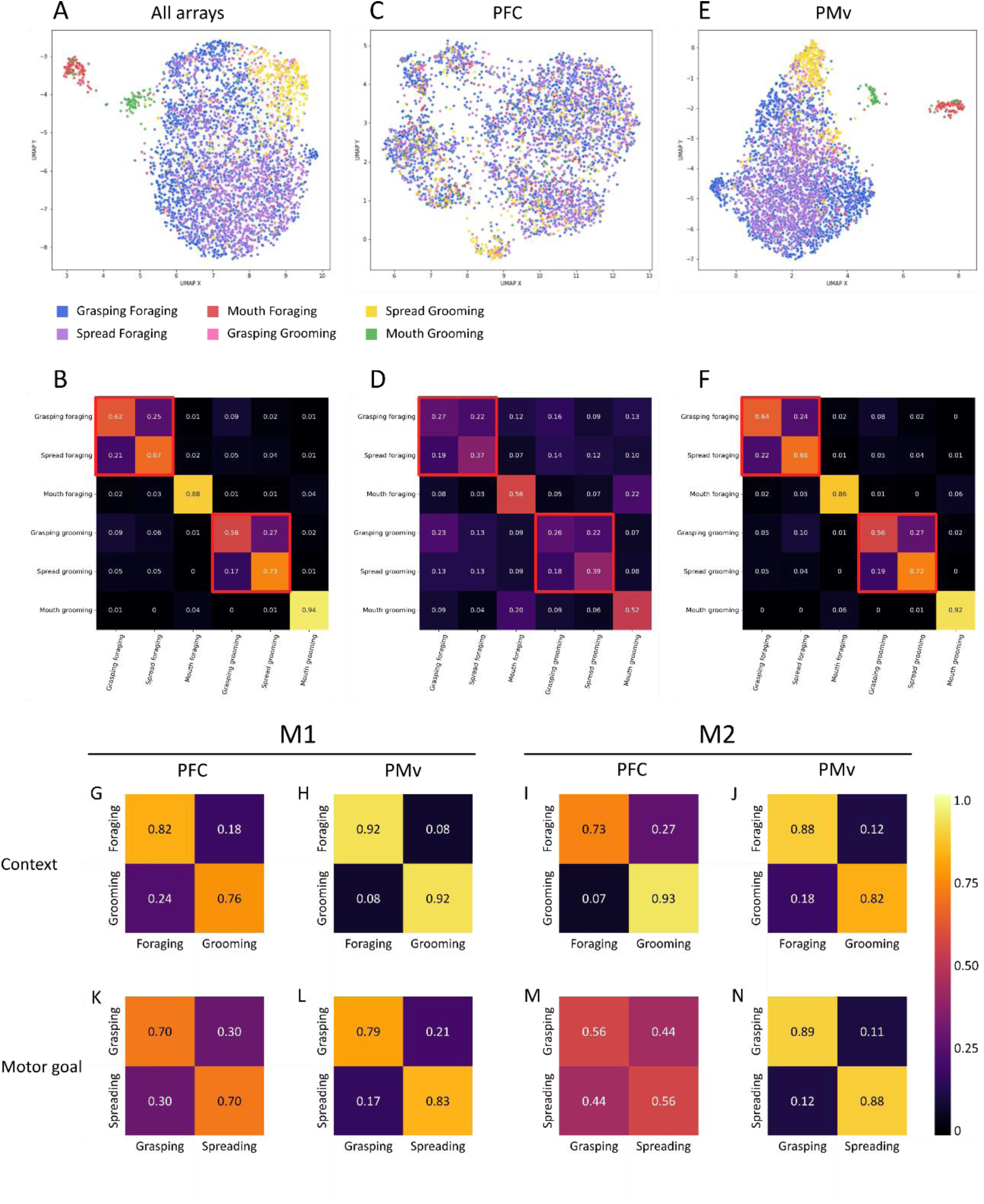
Population-level classification of behaviors in M1 using UMAP and XGBoost, and grouped classifier performances. (A, C, E) UMAP projections of vlPFC and PMv units (left), only vlPFC (middle) and only PMv (right), across all behaviors of all M1 sessions. (B, D, F) Confusion matrices of classifier performances trained on the same units as the respective UMAPs above. B) both areas confusion matrix, macro-F1 score = 0.647. Red boxes indicate overlap between hand grasping and hand spreading within each context. Mouth behaviors are well separated (mouth foraging = 0.86, mouth grooming = 0.92). D) PFC-only classifier confusion matrix (macro-F1 = 0.267). F) PMv-only classifier confusion matrix (macro-F1 = 0.654). (G-N) Confusion matrices of XGBoost classifier performance after regrouping behaviors by context (G-J, top row) or by motor goal (K-N, bottom row). Columns show M1 PFC (G, K), M1 PMv (H, L), M2 PFC (I, M), and M2 PMv (J, N). Mouth behaviors were excluded to allow consistent cross-animal comparison.

**Table 3:** Classification by ability to discriminate and preference between the two contexts (Prefer Foraging vs Prefer Grooming) on units detected as modulated in both contexts. Equal column refers to units that are modulated by both behaviors in different ways, but neither modulation is much stronger in magnitude than the other, while Not Discriminating are units that are modulated in a similar way between the two contexts according to CBPT.

| Subject | Region | Hand Grasping |  |  |  | Hand Spreading |  |  |  | Mouth Grasping |  |  |  |
| --- | --- | --- | --- | --- | --- | --- | --- | --- | --- | --- | --- | --- | --- |
|  |  | Discriminating |  |  | Not<br>Disc. | Discriminating |  |  | Not<br>Disc. | Discriminating |  |  | Not<br>Disc. |
|  |  | Pref.<br>For. | Pref.<br>Groom. | Equal |  | Pref.<br>For. | Pref.<br>Groom. | Equal |  | Pref.<br>For. | Pref.<br>Groom. | Equal |  |
| M1 | PMv | 7 | 3 | 2 | 0 | 8 | 6 | 1 | 0 | 11 | 5 | 1 | 0 |
|  | vIPFC | 4 | 1 | 1 | 2 | 10 | 1 | 0 | 0 | 2 | 3 | 1 | 1 |
| M2 | PMv | 21 | 5 | 6 | 1 | 15 | 7 | 4 | 6 |  |  |  |  |
|  | vIPFC | 0 | 2 | 0 | 2 | 1 | 1 | 0 | 1 |  |  |  |  |

### Neurons exhibit grooming-driven modulation patterns

A group of neurons we identified exhibited marked modulation patterns for grooming behaviors over their foraging counterparts (e.g. Unit 41 in Fig. 2). This selective enhancement for grooming suggests the existence of “social neurons” in frontal cortex, i.e., neurons preferentially coding actions with a distinctly social function. 20 out of 105 units (19.05%, pooled across monkey) were grooming selective (i.e. had the strongest modulation for one of the behaviors in the grooming context, like Unit 67). In addition, 15 of these units (14.29% of 105 total units, pooled across monkeys) showed modulation according to all three motor goals (i.e. Hand Grasping, Hand Spreading, Mouth Grasping) in the grooming condition. These results show evidence that a population of neurons in the frontal cortex is encoding preferentially social behaviors because of their social goals, rather than motor features.

### Population decoding highlights stronger contextual and motor-goal encoding in PMv than PFC

To assess population-level encoding of behaviors, we applied UMAP for dimensionality reduction and trained an XGBoost classifier on the embedded neural activity. In M1 (Fig. 4A), where units from both PMv and vlPFC were combined, the classifier achieved a macro-averaged F1 score of 0.647. On the same indicator, the classifier for M2 reached a score of 0.811 (results of M2 reported in Supp. Mat., Fig. S4). The UMAP embedding (left panel) shows a clear separation between the two contexts (foraging vs. grooming), while the two hand motor goals—grasping and spreading—largely overlap within each context. By contrast, mouth behaviors form distinct and well-separated clusters, reflecting strong differences in encoding between hand and mouth effectors. This effect is especially evident in PMv, where mouth foraging and mouth grooming were classified with 0.86 and 0.92 accuracy, respectively. The confusion matrix (right panel) highlights this structure, with red boxes indicating the main overlap between grasping and spreading within the same context. To allow a consistent comparison across animals, mouth behaviors were excluded from the grouped-class analyses presented in Figure 4.

When the areas were analyzed separately, performance diverged strongly. Using only vlPFC units (Fig. 4B), classification dropped to a macro-F1 of 0.267, whereas PMv alone (Fig. 4C) reached 0.654, nearly identical to the combined result. Consistently with this pattern, M2 scores of vlPFC and PMv classifiers scored a macro-F1 of 0.495 and 0.758, respectively (Supp. Mat., Fig. S4). This indicates that premotor cortex drives the classification both because of its higher unit count and because its neurons carry clearer information about motor goals. Even so, most classification errors in PMv also reflected confusion between grasping and spreading within the same context, reinforcing the idea that context information may be more robustly encoded than fine motor-goal distinctions.

To compare directly the contribution of context versus motor goal, we regrouped the behavioral classes and retrained the classifier (Fig. 4G-N). The key result is that classification by context consistently outperformed classification by motor goal, particularly in prefrontal cortex. In M2 PFC the performance drop was very pronounced, with context yielding 0.839 and motor goal only 0.562. For M1 PFC, the macro-F1 scores for social context and motor goal seemed very similar (0.692 vs. 0.698), but this similarity was misleading: the weighted F1 scores revealed the same effect as in M1 (0.837 vs. 0.703). This apparent discrepancy is due to the disproportionate size of the classes of foraging and grooming. While only a small portion of foraging events were labeled as grooming by context classifier (18%), in absolute numbers this was larger than the size of the entire grooming class, skewing the precision metric disproportionately. This was only an issue in this instance, while in all other classifiers the different performance metrics were very consistent. Premotor cortex, on the other hand, encoded both dimensions robustly. In M1 PMv, F1 scores reached 0.853 for context and 0.807 for motor goal, while in M2 PMv they were 0.836 and 0.888, respectively. This indicates that PMv provides strong representations of both social context and motor goals, whereas PFC is more specialized for contextual encoding.

### Comparing wireless recordings with head-restrained conditions

We compared the neuronal modulation associated with grasping behavior during naturalistic foraging in the current wireless recordings against previously collected head-fixed data obtained with the same chronic implants during a structured, trial-based grasping task. To assess the correspondence between these datasets, we quantified, for each monkey, the number of recording channels in the premotor cortex containing at least one unit significantly modulated for grasping (data from vlPFC under restrained condition have not been analyzed).

In the head-fixed condition, modulated activity was observed on 30 of 64 channels (46.9%) in M1 and 39 of 64 channels (60.9%) in M2. In the wireless foraging condition, the proportions were highly comparable, with 31 of 64 channels (48.4%) showing modulation in both monkeys (Supp. Mat., Fig. S5). These results indicate a strong consistency in the spatial distribution of grasping-related neuronal activity across experimental contexts and recording modalities, suggesting that the underlying sensorimotor representations remained stable over time despite differences in behavioral constraints and recording configuration.

## Discussion

Current findings provide evidence that neural representations of action in freely moving macaques extend beyond motor dynamics and immediate goals to incorporate the social context in which actions occur. By recording simultaneously from the PMv and the vlPFC during spontaneous naturalistic behaviors, we identified units that selectively encode socially directed goals (Fig. 3). This expands upon the traditional view of PMv coding motor acts only (*1*, *11–13*, *15*) by revealing its capacity to integrate the social relevance of actions, such as grooming a conspecific, into goal-directed neural codes.

The central result of this work is that a number of neurons in both the PMv and the vlPFC reliably discriminate between social and non-social behaviors. Importantly, these distinctions persist even when the patterns of motor acts used are very similar, indicating that neural coding reflects the social meaning of the action rather than just the motor goal. Although the free-behavior setup offers limited control over which behaviors occur, our design, focusing on four spontaneous natural behaviors categorized according to the motor goal (grasping vs spreading) and the social context (foraging vs grooming), enabled meaningful comparisons of naturalistic actions and revealed which aspects were most strongly encoded in the neural signal (Fig. 4G-N). For example, grasping food from the ground and grasping particles from a partner’s fur share the same motor goal (grasping), and the same or similar type of motor act (precision grip), yet their neural representations diverge according to whether they occur in a foraging or grooming context (Fig. 4A-F). This suggests the presence of dedicated neural subpopulations tuned to the social dimension of the action, supporting the idea that both premotor and prefrontal regions integrate motor plans with higher-order social context information (Table 4).

**Table 4:** Units selective or modulated by grooming. Selective: number and percentage of units selective for grooming across the six point-event behaviors (i.e. had the strongest modulation for one of the grooming behaviors). Modulated: number and percentage of units selective for grooming and modulated during all three grooming behaviors (i.e. hand, mouth, and spread grooming).

| Monkey | Area | Total units | Grooming Selective (n%) | Grooming Modulated (n%) |
| --- | --- | --- | --- | --- |
| M1 | PMv | 37 | 9 (24.32%) | 8 (21.62%) |
|  | vIPFC | 11 | 2 (18.18%) | 2 (18.18%) |
|  | Both | 48 | 11 (22.92%) | 10 (20.83%) |
| M2 | PMv | 42 | 6 (14.29%) | 4 (9.52%) |
|  | vIPFC | 15 | 3 (20.00%) | 1 (6.67%) |
|  | Both | 57 | 9 (15.79%) | 5 (8.77%) |
| Total | PMv | 79 | 15 (18.99%) | 12 (15.19%) |
|  | vIPFC | 26 | 5 (19.23%) | 3 (11.54%) |
|  | Both | 105 | 20 (19.05%) | 15 (14.29%) |

In line with recent work by Lanzarini and colleagues (*29*), our data revealed a high degree of mixed selectivity in both recorded areas. Many putative units were modulated across multiple, and in some cases nearly all, observed behavioral classes, often in distinct ways depending on the context (Table S2 and Fig. 2). Neurons dynamically changed firing patterns according to behavioral conditions, supporting the view that action goals, social cues, and motor plans are co-represented within a flexible and integrated neural code. This supports the idea of a shared representational format, where neurons encode dynamic motor synergies rather than discrete, goal-specific actions. Our results also align with the finding that motor coding in free-moving conditions diverges from that observed in restrained paradigms, revealing richer and more distributed representations (*29*). Such dynamic, context-dependent coding may represent an evolutionarily adaptive mechanism for supporting complex social behaviors in natural environments. Notably, most units that differentiated the same motor action across grooming and foraging contexts showed stronger modulation during foraging (Table 3). Combined with the larger share of the population encoding foraging relative to grooming, this pattern may reflect an evolutionarily privileged status of feeding and its neural embedding, which has been described as a very well-established motor synergy across both evolution and development (*33–35*). Further strengthening this point, we found also that, across both monkeys, the population in PMv, unlike vlPFC, had a significantly higher firing rate during goal-directed behaviors such as foraging compared to non-goal-directed ones (Fig. 1D). This result falls perfectly in line with prior literature about the role of the premotor cortex in planning and execution of goal-directed actions (*36*, *37*, *21*), and it shows how the neuroethological approach can increase the ecological validity of prior results by confirming them in the more naturalistic freely-moving setting. Indeed, despite the weaker modulation of the activity in vlPFC, our results also show the activation of some of its units during specific motor acts (i.e. hand grasping), in line with previous work that showed the importance of this area for the encoding of goal-directed actions (*16–18*).

Albeit small, the population of units we found in vlPFC responding during state-based grooming behavior showed predominantly negative modulation compared to resting baseline (Fig. 1E), consistent on both monkeys. Despite small methodological differences, this finding is in line with recent work of Testard and colleagues (*30*), who reported that the vast majority of putative units in vlPFC had a reduced firing rate during grooming behaviors. Notably, our dataset goes beyond state-level analysis. By differentiating sustained grooming epochs from discrete motor events, we show that many of the same neurons that decrease firing during the grooming state exhibit increased activity during specific hand and mouth actions embedded within the grooming bouts (Table 2, 4). This pattern reveals how neurons in vlPFC can dynamically change levels of activity to reflect contextual information and maintain their tuning to goal-directed actions like their premotor counterparts.

The population-level analysis we conducted also revealed striking structure in neural state space in both monkeys, reflecting the richness of encoding in spontaneous behaviors (Fig. 4A-F). The pattern of performances of the classifiers run on different datasets and groupings showed which behavioral and contextual features were more strongly reflected in the neural code. The population of units in PMv encoded clearly both the motor goal and the social context of the behaviors of both monkeys. Both the UMAP space embedding and the pattern of the confusion matrix indicated that the main drive of separation between behaviors was the social context. This result provides strong support for the involvement of PMv in integrating social information in motor planning. In contrast, prefrontal areas showed relatively weaker performance for the motor goal split, consistent with prior findings that vlPFC, being more distant from motor output, encodes motor details less strongly than PMv (Fig. 4G-N).

While freely moving paradigms open new windows onto naturalistic neural dynamics, they also require new analytic and experimental strategies. In order to address the absence of defined baselines, we developed a three-pronged method composed of the entropy-based metric of modulation (Supp. Mat., Fig. S5), the CBPT and the response profile. This approach proved reliable compared to standard GLM testing, and could serve as a general framework for analyzing neural activity in neuroethological settings, while still allowing meaningful comparison to previous studies. Indeed, recordings from the same chronic implant in restrained conditions revealed a comparable proportion of PMv channels encoding hand grasping (Supp. Mat., Fig. S5), demonstrating the reliability of the wireless system and highlighting the added value of naturalistic paradigms, where the very same recording site now shows modulation not only for grasping, but also for socially meaningful actions like grooming. Another issue is the variability in spontaneous behavior across sessions and between subjects (Fig. 1C), which creates a significant challenge for the reproducibility of the results. We mitigated this limitation by concatenating the sessions to increase the data pool, while others have incorporated structured enrichments to promote reproducible behavioral patterns (*38*), showing how this is a shared concern in the new field of neuroethology.

Building on recent advances in wireless neurophysiology in free-moving conditions our work highlights a new and complementary role of the motor system in natural social interactions. While Testard and colleagues (*30*) uncovered widespread population coding of social signals in higher-order cortical areas (prefrontal and temporal cortex), and Lanzarini and collaborators (*29*) emphasized the increased complexity of motor representations in free behavior, our results show that specific neuronal populations in the frontal cortex are tuned to social action goals. This offers a bridge between motor control and social cognition. Our findings uniquely demonstrate that the premotor cortex, which is traditionally associated with sensorimotor transformations, is also dynamically engaged by social contexts. This suggests that premotor and prefrontal neurons contribute not only to the control of movement, as demonstrated in traditional neurophysiological investigations (*1*, *5*, *36*, *39*), but also to the flexible integration of social and environmental information, thus expanding current models of frontal lobe function.

In summary, our study provides direct evidence that a number of neurons in PMv and vlPFC are selectively modulated during social behaviors in freely moving primates. According to this new perspective, as well as underlying goal-directed action, the frontal cortex also serves as a key hub where movement and social meaning converge. Altogether, our findings underscore the importance of incorporating ethologically-relevant paradigms (i.e. freely moving animals) in neuroscientific investigations. As neuroscience enters more ecological terrain, we may need to revise classical assumptions about functional specialization and neuronal specific tuning, adopting a more integrated view of perception, action, and social behavior.

## Acknowledgments

The initial setup of the ECR was done by research engineers Lucas Maigre and Thomas Perret, who created the video recording camera system, curated the EthoLoop implementation and the devices necessary to synchronize video, audio and neural recordings.

We want to thank the staff of the housing facility of the monkeys, led by Fidji M. Francioly, who cared for the continuous well-being of the animals.

We want to thank Pierre-Aurelien Beuriat for the surgeries on the animals.

For the video analysis of the behavior of the animals, due to the complexity and size of the dataset and the continuous refinement over the years of the research question and the ethogram of behaviors to study, the following people aided at different steps with manual annotation of the video recordings: Lilas Robert, Marine Cuvilliez, Cristina Rotunno, Nicolò Spada, Silvia Cassani.

Cristina Rotunno participated in theoretical and methodological discussions about the article, and along with Holly Rayson and Silvia Spada, proof-read the manuscript and provided feedback to the authors.

The AI tool ChatGPT (GPT-5, OpenAI) was used only for improving the English language of the manuscript, as none of the authors are native English speakers, and for enhancing code readability with comments.

## Funding

University of Lyon 1 Claude Bernard, PhD funding grant (JBA)

LABEX CORTEX (ANR-11-LABX-0042/ANR-11-IDEX-0007) of Université de Lyon operated by the French National Research Agency (ANR) (PFF)

ANR, ANR-20-CE37-0015-03 (Freemonk) (JRD, PFF)

Fondation pour la Recherche Medicale, ECO202306017335 (GCA)

Fondation pour la Recherche Medicale, LS293654 (EDI)

European Research Council (ERC) under the European Union’s Horizon 2020 research and innovation program (grant agreement no. 885746) (JRD)

Fondation pour la Recherche sur le Cerveau & Rotary (Espoir en Tête program) (JRD, PFF)

## Author contributions

According to CRediT model

Conceptualization: JBA, EDI, ACM, GCA, GCO, PFF

Data curation: JBA, EDI, ACM, GCA, GCO

Formal analysis: JBA, EDI, ACM, MBI, GCO

Funding acquisition: JRD, PFF

Investigation: JBA, EDI, GAN, PFF

Methodology: JBA, EDI, ACM, GCA, MBI, GCO, PFF

Project administration: JBA, JRD, PFF

Resources: MBI, JRD, PFF

Software: JBA, ACM, GCA, MBI

Supervision: JBA, MBI, GCO, PFF

Validation: JBA, EDI, ACM, GCA, MBI, GCO, PFF

Visualization: JBA, ACM, GCA, GCO, PFF

Writing – original draft: JBA, ACM, GCA, PFF

Writing – review & editing: JBA, EDI, ACM, GCA, MBI, GCO, JRD, PFF

## Competing interests

Authors declare that they have no competing interests.

## Diversity, equity, ethics, and inclusion

One of the authors identifies as a member of the LGBTQ+ community. This aspect of identity did not influence the study design or analysis.

## Data and materials availability

All data supporting the findings of this study will be made available upon request and deposited in a public repository such as Dryad upon acceptance for publication. Code used for analyses will also be made available. There are no restrictions on data availability.

## Materials and Methods

### Animals

Four adult female rhesus macaques (*Macaca mulatta*) participated in our experiment, with each pair consisting of one monkey implanted with electrodes and one unimplanted partner. The two implanted monkeys (M1 and M2) were 7 and 15 years old, respectively. Each implanted monkey lived with its respective partner (pM1 of age 7 and pM2 of age 11) in their own home enclosure for more than two years and thus had a well-established social hierarchy and dynamics. The determination of which monkey was the dominant in the relationship was based on multiple behavioral markers such as: amount of grooming received vs given (i.e. subordinate monkeys usually perform more grooming on the dominant), feeding priority in the cage when given food by the experimenters (normally the dominant would be at the forefront and the other would wait her turn), and aggressive behaviors such as mounting, threatening displays and chasing (40). All the procedures of the experimental protocol, as well as the housing and handling of the animals, complied with the European guideline (2010/63/UE) and were authorized by the French Ministry for Higher Education and Research (APAFIS #35051-2022013110567743 v5). The facility provided an enriched environment for the macaques.

### Surgical Procedures for Electrode Implantation

All surgical and experimental procedures were performed in compliance with European guidelines and the French National Committee in charge of the care and use of laboratory animals; they were also reviewed and approved by the ethics committee CELYNE (Comité d’éthique Lyonnais pour les neurosciences expérimentales, C2EA 42).

Surgeries were performed by neurosurgeon Pierre-Aurelien Beuriat and senior research investigators and Pier Francesco Ferrari, assisted by veterinarian anesthesiologists. Surgical plans were prepared using brain MR images from a Siemens PRISMA 3T scanner. T1-weighted images (MP-RAGE) were obtained to reconstruct the 3D cortical surface and vasculature. All surgeries were performed under isoflurane anesthesia and strict sterile conditions.

The surgical procedure, adapted from Fluet and Baumann (11, 41), involved implanting six floating multielectrode arrays (FMA, Microprobes for LifeScience, Gaithersburg, MD, USA, Supp. Mat., Fig. S1G, H)) in different cortical areas of the right hemisphere for monkey 1 (M1). For monkey 2 (M2), five FMAs were implanted in the left hemisphere, as shown in Figure 1B. In addition to the arrays, a titanium head post was fixed on the head of the animal. For a month after the surgery, the monkeys were treated with analgesic and anti-epileptic drugs to minimize discomfort and risks of damage to the cortex.

Each array consisted of 32 platinum-iridium electrodes with lengths between 1 and 6 mm and an inter-electrode distance of 400 µm. The electrodes had an impedance of 0.5 MΩ at 1 kHz. The arrays were implanted in the positions represented in Figure 1: one in the primary motor cortex (F1), two in the ventral premotor cortex (F5 hand and mouth representations), one in the dorsal premotor cortex (F2), and two in the prefrontal cortex (45a and 46/12r). This study presents data related to three primary cortical frontal regions: the ventral premotor cortex (F5 – two arrays) and the prefrontal areas 45a and 46/12r.

### Wireless Recording Device

The device used to perform the wireless recordings was the NeuroLogger made by Deuteron Technologies (RatLog-128, Deuteron technologies, Jerusalem, Israel). This device can be easily connected to the electrode recording system implanted on the head of the monkey, and is contained inside a customized 3D-printed PLA (polylactic acid) box designed to keep it safe from potential damage caused by manipulation. The antenna of the NeuroLogger communicated with a receiver connected to a computer to synchronize the electrophysiological recording with the behavioral events recorded during the tasks. The logger amplified and digitized the signal from the electrodes and stored the data on a memory card locally, from which they can be retrieved after the recording session ended. Connecting the device to the monkey’s head takes approximately five minutes, minimizing discomfort by reducing the duration of restraint. Once connected, the only constraint on the duration of the recording session was the power supply of the NeuroLogger, which was powered by a lithium battery (3.7 V, 250-350 mAh) providing approximately three hours of autonomy. During this period, the animal was free to move without the risk of damaging the recording system or causing injury.

### Experimental Setup during Free-Moving Social Interactions

The EthoControl room (ECR) is a cage measuring 2.25 x 1.94 x 2.11 meters (length x width x height), constructed with one-centimeter thick transparent anti-reflex glass walls, providing clear visibility from all angles. During the training period before the data acquisition reported in the current work, all monkeys had several hours-long habituation sessions in the EthoControl room over multiple months to make them feel more comfortable, free to explore the 3D space and produce their natural behaviors. The ECR was enriched with a wooden bedding material, rope for climbing, and to promote exploration and free roaming in the cage, we scattered small pieces of fresh and dried fruits on the floor to entice food search and foraging behavior. The experimental session and data acquisition began with four key steps: 1) The implanted monkey was first transferred from her home enclosure to a primate chair where the neural logger device was installed. 2) Once the recording system was secured, the monkey was placed in a transfer cage and transported to the ECR. 3) Neural and video recordings were initiated, and the monkey was released into the ECR to begin roaming and foraging. 4) Finally, the second monkey of the pair was moved from her home enclosure to the transfer cage, brought to the ECR, and released to join the partner. At the end of the recording session, the same procedure would be repeated backwards. The monkeys were trained so that when the experimenters would open the door of the transfer cage the partner monkey would go in first and be taken back to the home cage. Then the implanted monkey would go into the transfer cage, and from there back to the primate chair to remove the neurologger. After retrieving the recording system and closing the chronically implanted Omnetics connectors, the monkey would be taken back to the home cage and the data would be transferred from the SD card of the Neurologger.

Inside the ECR, perches and grooming boards were installed with the specific aim of encouraging a variety of motor behaviors during foraging. These enrichments provided opportunities to interact with objects and tools that were not accessible in the animals’ home cages, thereby stimulating more complex and varied activity patterns. During the recording sessions, the door of the ECR was kept closed, to avoid visual contact between the subject monkeys and other monkeys in the animal facility. Eight fixed Ximea MQ013CG-ON-S7 cameras were strategically positioned around the cage and connected to a synchronized acquisition system. Each camera was interfaced with a dedicated NVIDIA Jetson unit (equipped with 500GB SSD and WiFi), enabling precise temporal coordination of multi-angle video recordings.

EthoLoop, an object tracking software (31), utilized images captured by these cameras to track the positions of the monkeys’ colored collars within the cage, calculating each monkey’s coordinates in real-time. The positional data for each monkey was then transmitted to an automated motorized device that adjusted the orientation of two close-up cameras (one for each monkey) to continuously follow and record their behavior as they moved around the cage. The cameras used were the same model as the fixed ones, with the addition of adjustable focus lenses to maintain perfect video quality while following the monkey at different distances in the cage (lens model: Optotune −2 to +3 Diopter, M27 x 0.5 to M40.5 x 0.5 Focus Tunable Lens | EL-16-40-TC-VIS-5D-M27; controller model: Gardasoft TR-CL180 Industrial Lens Controller).

### Behavioral Scoring with BORIS

To identify and extract the behavioral sequences of interest, we employed BORIS, an open-source software for ethological annotation (42). We defined an ethogram of ethologically relevant behaviors that our monkeys spontaneously produced in their home enclosures, mainly foraging and social behaviors such as grooming and lip-smacking. We divided state behaviors (e.g. Locomotion, Grooming, and Foraging) from single-point events, such as object-contact during a grasping movement, to study the related neural activity patterns in greater detail. To account for the intrinsic variability of natural behaviors and movements, we extracted as much information as possible from the video recordings. Each behavior was labeled with modifiers specifying features such as the hand performing the grasping or the substrate on which the monkey was foraging, and this information was later used in the analysis to determine if they played a significant role in neural activity. All the labeling of natural behaviors was done manually by two trained experimenters and then reviewed for confirmation. The labeling of single-point events involved reviewing the video recorded sessions frame-by-frame and recording the time-stamp of the frame best matching the desired behavior. For grasping and spreading actions, the first frame of contact between the monkey’s fingertips and the target. For state events, instead, we started each state on the first frame that would make the behavior recognizable, and ended it when a different state would start (see the table in the supplementary material for a description of each behavior).

### Definition of Behaviors

The ethogram of all behaviors produced by the monkeys was defined by experimenters with extensive experience in ethological observation and classification of behavior. For each behavioral state, operational heuristics were established to determine precise onset and offset times in the video recordings. A full breakdown of the total duration spent in each behavioral state is reported in Supplementary Table S3.

- Resting: the monkey is in an idle state, fully seated, and not engaged in any physical interaction with its social partner. Start: when it terminates an action and transitions into an idle posture without initiating a new activity. Stop: at the onset of a new behavior, marked by initiation of an action; minor positional adjustments do not constitute a transition to a new behavioral state.
- Locomotion: the monkey moves from one location to another within the cage or displays repetitive, stereotyped back-and-forth movement patterns. Climbing up and down from the platform is included. Start: when it initiates a continuous movement aimed at relocating within the enclosure. Stop: when it reaches a new position and either transitions into an idle state or initiates a new goal-directed behavior.
- Foraging: active food-seeking behavior, which may involve locomotion with active search and spreading bedding with the hand, or stationary manipulation with the hand of foraging devices (usually involving spreading and grasping with whole hand and/or precision grip). Start: when it directs its gaze and initiates purposeful interaction with the ground or a foraging apparatus. Stop: when it ceases searching and hand interaction with the ground or device.
- Eating: ingestion of food items outside of the foraging context. The monkey brings previously retrieved food (typically stored in cheek pouches) to the mouth and chews it. Start: when visible hand-to-mouth movement or chewing of stored food begins. Stop: when ingestion ceases and no further chewing or food handling occurs.
- Mouthing: oral exploration of non-food objects or surfaces, such as licking or chewing parts of the cage or other environmental elements. Start: when the mouth makes purposeful contact with an inedible object, accompanied by oral manipulation. Stop: when the oral contact is broken and the animal’s attention or activity shifts away from the explored item.
- Exploration: visually or manually investigating the environment, typically involving short repositioning movements to obtain better access or views of an object or area. Lifting and extending the body to explore a target at a higher location (hands are leaning or holding a substrate). This state is characterized by a sustained, focused gaze on the target. Start: when the monkey orients its gaze and initiates purposeful inspection of an object or part of the cage. Stop: when it disengages visual attention or ceases the exploratory actions.
- Grooming (allogrooming): the monkey engages in grooming behavior by manually or orally manipulating the partner’s fur or skin, involving actions such as parting the hair, removing particles with a precision grip, or inspecting specific areas. Start: when it directs attention toward the partner and initiates tactile exploration of the partner’s body. Stop: when it interrupts physical contact and disengages attention for at least one continuous second.
- Self-grooming: manipulation of the monkey’s own fur or skin using hands or mouth, including actions such as precision grip and pulling the fur, single bouts of scratching (unlike Scratching as a behavioral class, which is more repetitive and sustained), rubbing, or inspecting specific body areas. Start: same as Grooming but directed to Self. Stop: same as Grooming.
- Received grooming: the monkey receives grooming from the partner. Start: when the partner initiates grooming contact. Stop: when the partner interrupts the grooming bout.
- Scratching: repetitive scratching movements directed at the monkey’s own body using one or both hands. This behavior can occur spontaneously or during transitions between other states. Start: when the hand begins a repetitive scratching motion on the body surface. Stop: when scratching movements cease for at least one continuous second.
- Lying: the monkey adopts a fully or partially horizontal posture, usually on a flat surface such as the cage platform. Lying often occurs in the presence of the partner and is frequently followed by Received Grooming. Start: when the animal settles into a horizontal resting position with trunk and limbs supported by the surface. Stop: when it raises the torso or initiates another active behavior.
- Presenting hindquarters: the monkey changes posture to turn away and present the hindquarters or the back to the partner. Start: when the animal orients toward the partner and presents a body region while maintaining a still, expectant posture. Stop: when the posture is discontinued or when grooming begins.
- Lipsmacking: rapid rhythmic movements of the lips, typically associated with affiliative or appeasement signals. This can be directed toward the partner, the experimenter (often off-camera), or occasionally toward other monkeys visible or audible outside the room, especially at session beginnings or endings. Start: when repetitive lip movements characteristic of lipsmacking begin. Stop: when these movements cease for at least one second or are replaced by another behavior.

For the scope of the analysis reported in Fig. 1E between goal-directed and non-goal-directed, we divided the groups according to the following criterion. Behaviors were assigned to goal-directed if they included motor acts with hand of mouth effectors and involved an intentional goal. Non-goal-directed were considered the behaviors that involved repetitive or automated motor acts, such as Locomotion, or that did not involve motor acts with hand or mouth effectors, such as Exploration and Lying. From this grouping we excluded Scratching because it is a topic of debate whether it is a goal-directed action or not, and Resting because it was defined as the absence of other behaviors and is more of a baseline (32).

In a similar fashion, we created an ethogram for the point events to label specific movements of the monkeys and to allow later to align those behaviors for a trial-based analysis. The frame of the video on which to label each movement was selected by the experimenters, to the best of their abilities, according to the definition provided here. The first visible frame of contact between the effector (i.e. hand or mouth) and the target (i.e. bedding, food or fur) was marked with the event. Crucially, while it was not explicitly measured, all of the point events coded required the head of the monkey to be directed towards the target of the action, as they are precise goal-directed movements that require focused visual attention. This minimized the variability and impact of gaze behavior on the neuronal activity during grooming and foraging behaviors. A full breakdown of the number of behaviors produced by each monkey is reported in Supplementary Table S2.

- Grasping: the monkey performs a goal-related manual action involving the use of the hand to reach out for or hold an item. This may include retrieving food items or objects from the partner’s fur during grooming interactions, or from the substrate or a foraging board during foraging behavior
- Spreading: the monkey uses its hand (mainly the fingers) to part or displace materials in a deliberate manner. This includes separating the partner’s fur to access the skin during grooming, or manipulating substrates such as wood chips or the bristles of a foraging device to locate concealed food items during foraging. It usually involves lateral movement of the distal part of the arm
- Mouth grasping: the monkey positions its body to directly access the partner using the mouth (with a lips protrusion, called “pout face”), typically leaning forward with the torso to inspect or manipulate the fur or skin without the use of the hands. During foraging, the monkey lowers its head toward the ground while flexing the forearm and extending the lips to directly retrieve a food item without using the hands

### Neural and behavioral recording and processing

We recorded electrophysiological extracellular activity from a total of 128 channels from each monkey. Two chronic multi-electrode arrays of 32 channels each were implanted in the prefrontal cortex and two in the premotor cortex. The electrodes were connected to the neural datalogger system from *Deuteron Tech* through Omnetics connectors and the signal was stored locally in the SD card of the system. At the beginning of a recording session, we would use the LoggerCommand3 Deuteron software to communicate with the device and synchronize the time of the recording. After launching the recording, we would start the EthoLoop software for controlling the camera system and record the behavior. At the same time as the start of the video recording, a TTL signal was sent by EthoLoop to the transceiver which communicates with the neural logging device. This signal was then used during the analysis to align the behavioral recording to the neural recording.

The recording was amplified, digitized, bandpass filtered (2-6000 Hertz) and then stored locally on the recording device. After the session was concluded and the device was retrieved from the monkey, the data would then be downloaded from the SD card and transferred to permanent storage for future analysis.

### Spike sorting

To analyze the neural data, we used the spike sorting framework *SpikeInterface ver. 0.97.1* (45) to develop our own pipeline for preprocessing and spike sorting the signal. The first step of the pipeline was bandpass filtering (300-6000Hz), followed by whitening of the recording, notch filtering, common reference subtraction (global reference), peak detection with a threshold of 3.5 standard deviation from the mean signal of each channel, and ultimately spike sorting with the integrated module to run the *PyKilosort* algorithm (43) on each channel to extract sorted units.

The curation was performed manually with the standard *Phy* software (44) for visualization and curation of spike sorting outputs. First, we removed artifacts of the recording, then removed all units that showed a clearly non-physiological waveform. The remaining putative units were then subdivided into multi-units and single-units. In order to select well-isolated single units from the remaining pool we used two metrics implemented in the SpikeInterface framework: inter-spike intervals violations ratio (ISI) and signal-to-noise ratio (SNR). The first quantifies the level of contamination of the unit by identifying the spikes that cannot possibly be from the same unit because the interval from the previous spike is too short to be physiologically plausible. The computation of the ISI violations is implemented as described in (46), and the threshold used was 0.25, where any unit with a lower level of contamination was considered well-isolated. The second metric computes the ratio between the maximum amplitude of the average waveform of the unit and the standard deviation of the background noise on the same channel. A unit with a high SNR has a greater signal than the noise and is therefore more likely to correspond to a neuron (47, 48). For our analysis, we chose a threshold of 3 snr and excluded from the single unit group all the units below that number.

In addition to those hard thresholds, additional criteria taken into consideration were the presence ratio of the unit throughout the entire recording and the stability of the firing rate. Units that were exhibiting inconsistent patterns (i.e. clearly visible variations of firing rate from session to the other) were excluded from the analysis. We also merged units that were located on the same channel and showed the same average waveform, similar principal component analysis (PCA) components and similar amplitudes. Opposite to this procedure, in some cases the sorting algorithm clustered together spikes that were belonging to two different units, which could be easily identified examining the PCA plots on *Phy*, and the clusters could be cut manually to assign the spikes to two new clusters.

For the analysis described in this paper, both well-isolated (single-units) and non-well-isolated (multi-units) putative neurons found with this method were taken into consideration. To consider the three sessions of each monkey as a whole, we concatenated the data from each session consecutively and then applied the same method above. The spike sorting algorithm was able to identify units that appeared during all three recordings, albeit with variable degrees of precision. Some units were not found consistently across all three recordings, so they were manually excluded.

### Single-unit responses

After defining the pool of units to consider in the analysis, we aligned the behavioral events to the neural recordings by adding to the timestamp of the events the difference between the start of the neural recording and the start of video acquisition. The first method we used to compute the average firing rate of each unit was to divide the total number of spikes by the duration of the session. Second, we computed the firing rate during periods when the monkey was idle. We did so by considering only the time bins classified as Resting behavior, and counting the number of spikes divided by the duration of the Resting. The results of the first and second methods had negligible differences, so we used the first method for the subsequent analysis.

### Statistical analysis of firing rates across behavioral groups

To test whether the population’s neuronal activity differed between goal-directed and non-goal-directed behaviors, we performed a nonparametric permutation test on the average firing rates computed for each unit (using Matlab software). For each neuron, the mean firing rate was calculated separately for each behavioral state. Behaviors were then grouped into two categories: goal-directed (Foraging, Eating, Mouthing, Grooming, and Self-Grooming) and non-goal-directed (Exploration, Locomotion, and Lying). The observed difference in mean firing rates between the two groups was compared against a null distribution generated by 10,000 random permutations of group labels. The *p*-value was computed as the proportion of permutations yielding a difference greater than or equal to the observed one. This approach makes no assumptions about the underlying distributions of firing rates and provides robust inference at the population level.

### Measure of effect size of modulation against resting state

To quantify modulation of single-unit activity across behaviors, we compared firing rate distributions during each behavioral state with those during identified rest epochs using the Cohen’s D effect size. This metric, adopted from Testard et al. (30), provides a standardized measure of mean difference and enables direct comparison between the two studies.

For each monkey separately, and for each array (F5hand, F5mouth, 46v, 45a), we extracted firing rate distributions using 1 s bins as independent samples. Because behaviors occurred with variable durations, we downsampled each distribution to a maximum of 100 samples per state to equalize sample size across behaviors and between state and rest. When fewer than 100 samples were available, all observations were used. The same number of samples was randomly drawn from the rest distribution to match. This subsampling procedure was repeated 10 times to obtain stable estimates, and the resulting Cohen’s D and p-values were averaged across runs.

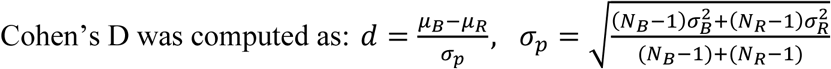

where *μ*_*B*_, *μ*_*R*_ and *σ*_*B*_, *σ*_*R*_ represent the means and standard deviations of the two distributions (behavior vs. rest), and N_B_, N_R_ their sample sizes. The resulting matrix of Cohen’s D values was plotted as a heatmap using a red–blue color scale, spanning from +1.5 to −1.5 (red means positive modulation compared to rest). Arrays were ordered vertically as premotor (F5hand, F5mouth) followed by prefrontal (46v, 45a), separated by a horizontal black line. A two-sample t-test was performed for each unit-by-behavior pair, yielding one p-value per comparison. Multiple comparisons were controlled using the Benjamini–Hochberg False Discovery Rate (FDR) correction with α = 0.01 applied across all tests. Units showing FDR-corrected p < 0.01 were considered significantly modulated.

### Shannon’s entropy metric

To overcome the lack of baseline in freely-moving recordings, we developed an entropy-based metric of firing patterns to assess whether a unit was significantly modulated during a given behavior. This method bypasses the need for a baseline and measures whether firing during a time window around an event is structured in a meaningful way. In order to assess whether a unit is responsive or not to a specific behavior of the monkey, we used a method based on Shannon’s information theory (49). The idea is that the amount of entropy in a system represents the amount of uncertainty in it. The higher the entropy, the more chaotic is the behavior of the system, while the lower the entropy, the higher is the amount of information. Entropy quantifies how evenly the probabilities of possible events are distributed within a system. When the probabilities are uniformly distributed, entropy is maximized, indicating greater uncertainty and less information. Conversely, when the distribution is highly skewed, meaning some events are much more probable than others, entropy is reduced, and the system carries more specific information. This is encapsulated in this mathematical formula, for which it can be found that the maximum entropy occurs when the probabilities follow the uniform distribution, and the null entropy is obtained when the probabilities follow the Dirac’s Delta function – where the probability is 1 for one value of x and 0 for all other values. The second case represents a scenario in which we know with absolute certainty when the event will occur, so we have all the information, while in the former case we have no information at all because the event is equally likely to occur for each value of x.

In our case, we modeled the firing of each unit during a window of time centered around the behavioral events as a distribution of probabilities of that unit emitting a spike at different times around the event. We convolved each spike train with a Gaussian kernel *σ* = 50*ms* to smooth the activity, and normalized the values to compute at each 1ms time point *t* over the 1000ms window *T* the probability of a spike occurring. We used the classic formula (ref Shannon) to compute the entropy *H*:

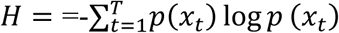

Where *x*_*t*_ is the event “occurring spike at time *t*” and *p*(*x*_*t*_) is the probability to observe the spike, which is intuitively the same thing as the instantaneous firing rate of the neuron. The underlying assumption is that when the activity of a neuron is influenced in any way by a task, that is reflected in a modulation of its firing rate, and therefore a distribution of probabilities more skewed, which means a lower entropy measure.

### Neuronal Specificity

We assessed whether each neuron was significantly modulated by specific behaviors by quantifying the uncertainty in its spike timing using Shannon’s entropy according to the process described in the previous paragraph. Lower entropy values indicate more structured, behaviorally modulated responses, whereas higher entropy reflects more random spiking activity. To determine whether a unit was task-related, we generated a null distribution by randomly permuting the spike times within the analysis window for each trial (100 permutations). We then computed the average entropy across all permutations, providing an estimate of the entropy expected under random, non-task-modulated activity. The entropy distributions from the original and randomized spike trains were compared using a Wilcoxon signed-rank test (MATLAB function *signrank*).

Because the number of trials per behavioral class varied, we subsampled the classes to match the smallest number of trials, repeated the random sampling and statistical testing 50 times, and averaged the resulting p-values to obtain a robust estimate for each unit. A neuron was considered significantly modulated for a behavior if its average *p-value* was *< 0.05*. Since each unit’s responses are considered independent, we did not apply a strict correction for multiple comparisons; however, when an FDR correction was applied, the significance of the results remained consistent.

### Neurons’ Polarity Analysis

After identifying behaviorally specific neurons via Shannon’s entropy, we then characterized the polarity of their responses by comparing each neuron’s behavior-aligned firing rate to its session-wide reference measure and measuring the signed area between them. Specifically, for each neuron-behavior pair, spike trains were aligned to the behavioral event (*–500 ms* to *+500 ms*), binned into contiguous *20 ms* windows, and converted to instantaneous firing rates (*Hz*) exactly as described above. We then computed the trial-averaged firing rate trace, *FR*(*t*_*i*_), and subtracted the neuron’s sessions-wide mean rate *FR*, calculated across all sessions, to obtain the deviation

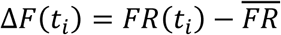

In the absence of a clearly defined pre-event baseline, *FR* serves as a stable, behavior-independent benchmark that reflects each cell’s intrinsic excitability without being skewed by behavior-related modulations.

We then quantified the magnitude and polarity of each neuron’s deviation trace *ΔF*(*t*) by numerical integration using the trapezoidal rule, which approximates the area under a sampled curve by dividing it into adjacent trapezoids

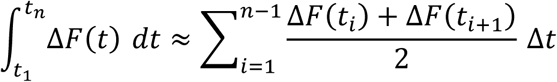

where Δ*t* = 20 ms is the bin width.

We then separated this total area into positive and negative contributions by summing only over bins where Δ*F* exceeded or fell below zero, respectively:

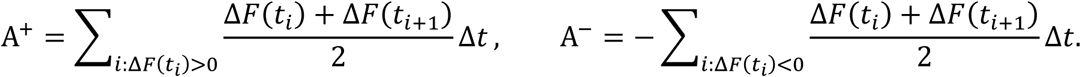

Each response was labeled facilitated if A^+^ exceeded A^−^, inhibited if A^−^ exceeded A^+^, and biphasic if both A^+^ and A^−^ were nonzero and each represented at least 10 % of the other’s magnitude.

- Facilitated: *A*^+^ > *A*^−^
- Inhibited: *A*^−^ > *A*^+^
- Biphasic: *A*^+^ ≠ 0, *A*^−^ ≠ 0, *A*^+^ > *A*^−^ ∗ 0.1, *A*^−^ > *A*^+^ ∗ 0.1

### Cluster-based permutation test

To investigate whether task-related neurons exhibited significantly different modulations between paired behavioral conditions (e.g., Grasping Grooming vs. Grasping Foraging), we conducted a cluster-based permutation test (CBPT, 50). We binned at 1ms the activity in a window of 1s centered around each behavioral event, as for the other analyses conducted in the paper. The analysis evaluated independent two-sample t-statistics at each 20ms time bin across trials independently for each neuron.

First, for each pair of modulated conditions of a neuron, the neural activity was normalized by z-scoring between both conditions. Subsequently, at each time bin, an independent two-sample t-test was conducted between conditions, yielding a series of observed t-values over time. Clusters were then identified as contiguous sequences of time bins where t-values exceeded the two-tailed critical threshold (*α = 0.05*). Positive clusters were defined as sequences of bins with t-values greater than the positive critical threshold, and negative clusters as sequences with t-values smaller than the negative critical threshold. For each observed cluster, the cluster mass was calculated as the sum of the t-values within that cluster. We then constructed empirical null distributions by performing random permutations (*n=1000*), in which trial labels were randomly reassigned across conditions. This resampling procedure disrupted any systematic association between neural activity and experimental condition, effectively simulating the null hypothesis of no difference between groups. For each permutation, independent two-sample t-tests were conducted at each time bin, followed by cluster identification using the same criteria applied to the observed data. Within each permuted dataset, positive and negative clusters were identified separately, and the mean cluster mass was calculated for each type. This process yielded empirical null distributions of mean cluster masses expected purely by chance.

Each observed cluster mass was then statistically evaluated by comparing it to the corresponding null distribution. Specifically, for each observed cluster mass, the p-value was calculated as follows:

For positive clusters:

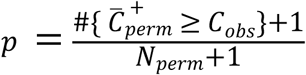

For negative clusters:

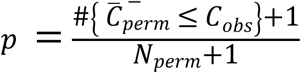

where 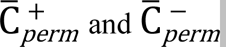 represent the mean positive and mean negative cluster masses obtained across permutations, respectively, and *N*_*perm*_ is the total number of permutations. #{·} denotes the count of events satisfying the condition inside the brackets. Clusters were considered statistically significant if their cluster-level p-values were below *α = 0.05*. By operating at the cluster level rather than at individual time bins, this method effectively controls the family-wise error rate (FWER) while maintaining sensitivity to sustained temporal differences between conditions. The CBPT was applied independently for each behavioral category (Hand Grasping, Hand Spreading, Mouth Grasping).

### GLM analysis of channel modulation in tethered recordings

To validate the entropy-based method, unit modulation in the tethered dataset was assessed using a generalized linear model (GLM) with a Poisson distribution, suitable for spike count data. For chair-restrained trials, the baseline of each trial was a period of *1s* of activity preceding the go-signal, during which the monkey is supposed to be waiting and not engaging in any other action, while the behavior is a 1s window of activity centered around the moment of contact between the fingers of the monkey and the target object of the grasping during the trial. Units with a significant condition coefficient were considered modulated. GLM results were compared with the entropy-based analysis to evaluate agreement and validate the baseline-independent approach for application to wireless recordings, which lack structured baseline periods.

### Temporal Processing of Neural Data

To characterize behavior-specific activation patterns, we treated the neural spike trains as temporal series. The processing pipeline consisted of the following steps:

1. **Temporal Windowing**: For each behavioral event, we extracted a 1-second window of neural activity centered around the event onset (−500 ms to +500 ms).
2. **Rebinning**: The original 1,000-element vector (1 ms bins across 1 second) was rebinned into a 62-element vector, with each bin representing approximately 16.13 ms of neural activity to match one bin of neural activity to one frame of video recording.
3. **Smoothing**: A Gaussian kernel with a bandwidth of σ = 2.48 was applied to smooth the rebinned vector, reducing noise while preserving the temporal structure of the neural activation patterns associated with specific behaviors. This smoothing step helps to create a more continuous representation of the firing rate modulation over time, suitable for pattern analysis.
4. **Normalization**: Each temporal series was detrended by removing the mean and scaling to unit standard variance to account for baseline differences in firing rates across units and recording sessions.

### Feature Extraction with MiniRocketMultivariate

To efficiently capture the patterns encoded in the neural temporal series, we employed MiniRocket Multivariate (*MINImally RandOm Convolutional Kernel Transform*)(51), a highly efficient multivariate time series feature extraction algorithm designed for time series classification tasks. It employs a fixed set of 84 convolutional kernels with pre-defined dilations and biases to transform input time series into discriminative features. MiniRocket optimizes kernel configurations deterministically, eliminating randomness while maintaining state-of-the-art accuracy. For each kernel, the algorithm computes the Proportion of Positive Values (PPV) pooling operator, which captures the fraction of positive activations in the convolved output, generating a compact yet expressive feature representation. This approach is extended to multivariate data by independently processing each dimension and concatenating features across channels, preserving inter-variable relationships while retaining computational efficiency.

The MiniRocketMultivariate algorithm was applied with the following parameters:

- Number of kernels per brain area= 5,000. Maximum kernel dilation= 32

Two separate feature matrices were generated, one for each brain area (Prefrontal and Premotor Cortex). These matrices were subsequently concatenated to create a unified representation of neural activity across both brain regions. The final feature matrix was standardized, with features detrended and scaled to unit variance.

We configured MiniRocketMultivariate to generate 5000 features (kernels) per input time series, with a maximum kernel dilation of 32. The transform was applied separately to the processed time series data from all units within the vlPFC and all units within the PMv for each trial. The resulting feature vectors from the two brain areas were then concatenated, creating a single, unified feature vector representing the combined neural activity across both regions for each behavioral trial. The final feature matrix was standardized, with features detrended and scaled to unit variance.

### Dimensionality Reduction for Visualization

For visual inspection and evaluation of the neural patterns, we reduced the high-dimensional feature space using Uniform Manifold Approximation and Projection (UMAP, 53).

UMAP is a non-linear dimensionality reduction technique that preserves both local and global structure in the data, making it well-suited for visualizing complex, high-dimensional neural representations. We projected the standardized feature matrix onto a 2-dimensional space using the following UMAP parameters: *n_neighbors=25, min_dist=0.1, and metric=’euclidean’*.

The resulting two-dimensional projections were used to build the UMAP representations of Figure 4A, C, E in the main text, and Figure S3A, C, E in Supplementary Materials

### Classification of Neural Patterns

To evaluate the extent to which the extracted neural features could discriminate between the different behavioral categories, we implemented a supervised machine learning classification pipeline.

### Data Partitioning

The feature dataset was divided into training and testing partitions using a stratified split, with 70% of samples allocated for training and 30% for testing. Stratification ensured that the relative proportions of behavioral categories were preserved in both partitions, which was particularly important given the imbalanced nature of our dataset. This approach maintained the same class distribution in both training and testing sets, providing a more reliable estimate of model performance across all behavioral categories. The training set was used for hyperparameter optimization, while the test set was initially used for a preliminary evaluation.

### Classification Model: XGBoost

We employed the XGBoost classification algorithm (52), an efficient implementation of gradient boosted decision trees.

Key configuration parameters for the XGBoost model included:

- objective: Set to ‘multi:softmax’ for the multi-class behavioral classification task.
- eval_metric: Set to [‘merror’, ‘mlogloss’] for monitoring model performance during training.
- tree_method: Set to ‘hist’ for faster computation.

### Hyperparameter Optimization

To find the optimal hyperparameters for the XGBoost model tailored to our dataset, we utilized a randomized search strategy. This search was performed exclusively on the training partition of the data. The process involved:

- Class Imbalance Handling: To address the imbalance in the number of samples per behavioral category, the training partition was resampled using a random under-sampling strategy prior to the hyperparameter search. This procedure balanced the dataset by reducing the number of samples in the majority classes to match the number of samples in the minority class, ensuring equal representation of all behavioral categories during model training. This resampling was applied exclusively to the training data partition.
- Cross-Validation: A 5-fold stratified cross-validation scheme was used within the search. The training data was split into 5 folds, maintaining class proportions in each fold. For each candidate parameter set, the model was trained on 4 folds and validated on the remaining fold, repeated 5 times.
- Parameter Search Space: The randomized search explored various combinations of hyperparameters drawn from predefined distributions. The following parameters and distributions were included:
- max_depth: Maximum tree depth, integer values sampled uniformly from [2, 5]
- min_child_weight: Minimum sum of instance weight needed in a child, integer values sampled uniformly from [1, 9]
- learning_rate: Step size shrinkage, values sampled from a log-uniform distribution between 10−3 and 0.5.
- n_estimators: Number of boosting rounds (trees), integer values sampled uniformly from [50, 399]
- gamma: Minimum loss reduction required to make a further partition, values sampled uniformly from [0, 1].
- subsample: Fraction of samples used per tree, values sampled uniformly from [0.6, 1.0].
- colsample_bytree: Fraction of features used per tree, values sampled uniformly from [0.6, 1.0].
- reg_alpha: L1 regularization term on weights, values sampled from a log-uniform distribution between 10−5 and 10.
- reg_lambda: L2 regularization term on weights, values sampled from a log-uniform distribution between 10−5 and 10.
- Class Imbalance Handling: To address the imbalance in the number of samples per behavioral category, class weights were employed during model training and optimization. These weights were calculated inversely proportional to the class frequencies in the training data, giving higher importance to minority classes.
- Search Iterations: The randomized search evaluated 100 different hyperparameter combinations (n_iter=100). With 5-fold cross-validation for each combination, this resulted in a total of 500 model fits during the optimization process.

### Final model and performance evaluation

After identifying the best set of hyperparameters from the randomized search (based on the average cross-validated performance, e.g., minimizing logloss or error rate), we estimated the final model performance using a separate 5-fold stratified cross-validation procedure applied to the entire training dataset.

This involved repeatedly training the model using the best hyperparameters on the training portion (4 folds) of each split and evaluating it on the remaining validation portion (1 fold). Crucially, within each cross-validation split, the 4 training folds were first balanced using the same random under-sampling technique described previously before the model was trained. This ensured that the model was trained on a class-balanced dataset in every iteration, while its performance was evaluated on the original, imbalanced distribution of the validation fold, providing a realistic estimate of its generalizability. Performance metrics were averaged across all folds to provide a comprehensive assessment of classification accuracy and generalizability (e.g. performance metrics in Fig 4).

**Fig. S1.**
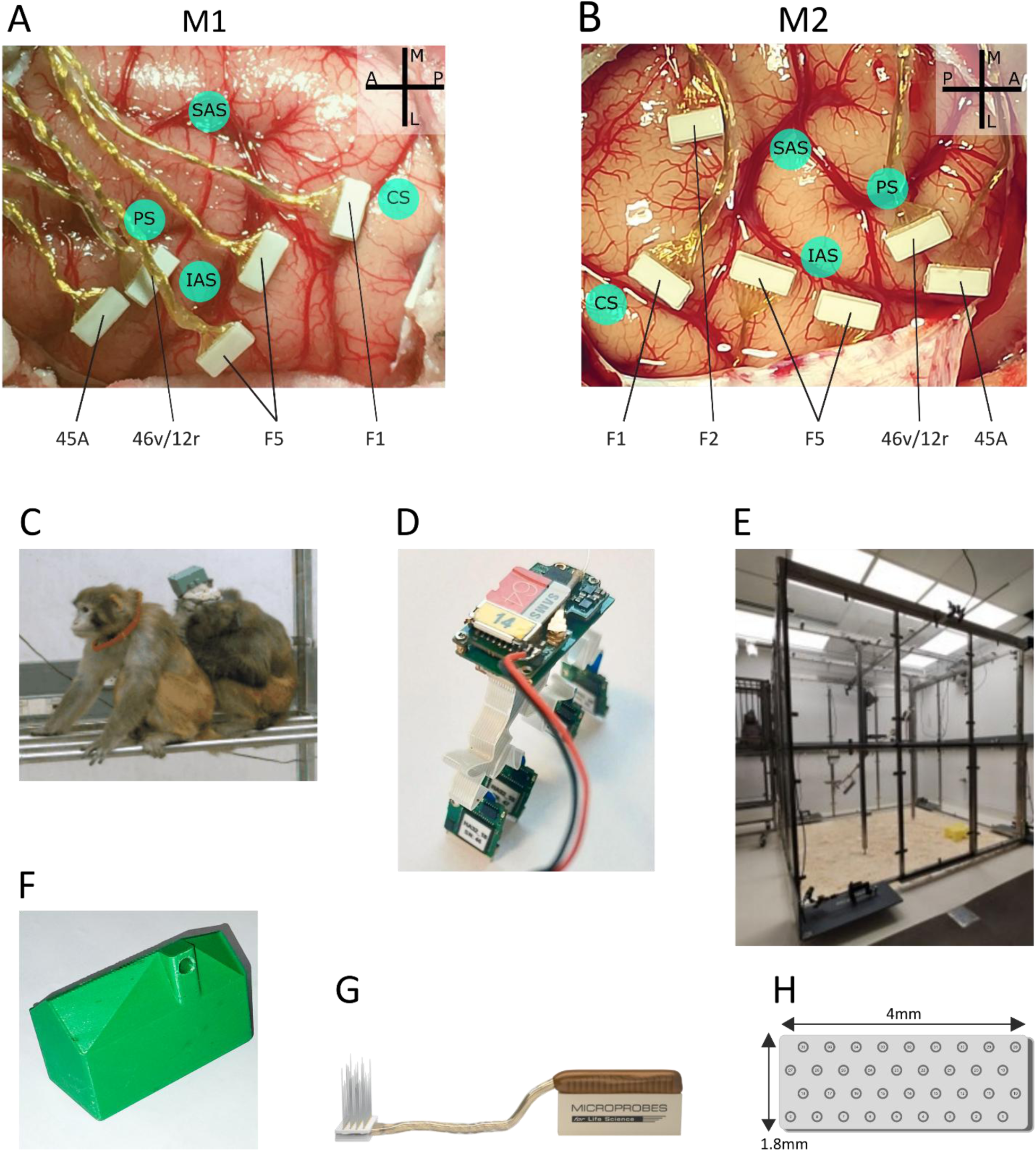
Chronic implant, EthoControl Room and Neurologger device. (A) Cortical implant in left hemisphere of M1. (B) Cortical implant in right hemisphere of M2. (C) Video frame exemplifying grooming behavior between M1 and her partner. (D) Neurologger device with custom-designed connectors for compatibility with the chronic implant. (E) Photograph of the EthoControl Room. (F) Photograph of the 3D-printed plastic box to secure and protect the Neurologger during the recording sessions. (G, H) Diagram of the MicroProbe FMAs implanted and geometry of the probe.

**Fig. S2.**
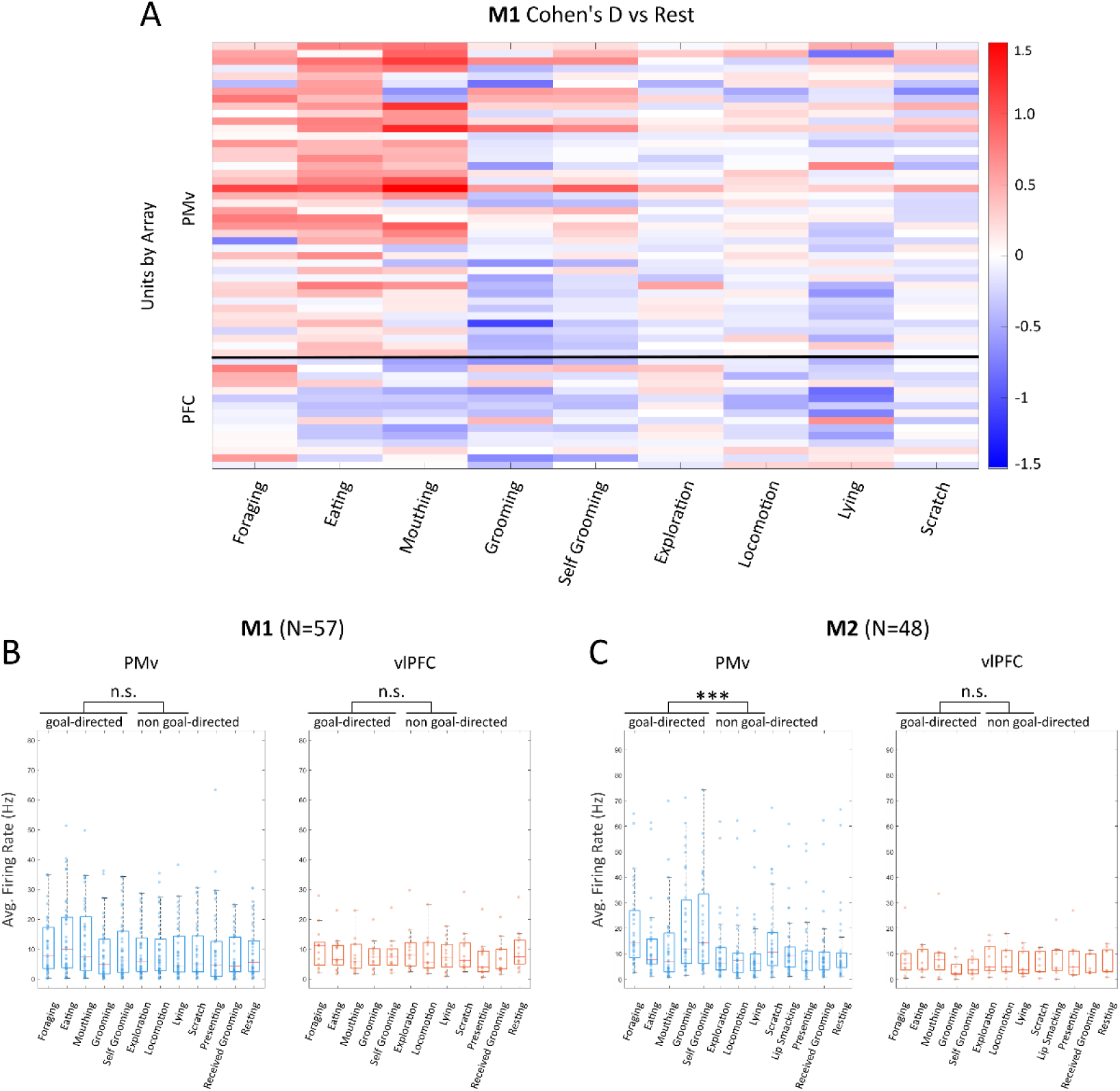
Firing rates boxplots and Cohen D heatmaps separate by monkey. (A) Cohen’s D figure of M1. (B, C) Boxplot of the same analysis performed separately for each monkey yielded qualitatively consistent results. For Monkey 2, PMv activity was significantly higher during goal-directed behaviors compared to non–goal-directed ones (p = 0.0005), whereas vlPFC showed no difference between the two groups (p = 0.945). For Monkey 1, the PMv difference did not reach the conventional significance threshold (p = 0.0736), but it showed a trend in the same direction, suggesting a similar underlying effect. In both monkeys, vlPFC activity was not significantly modulated by behavioral group.

**Fig. S3.**
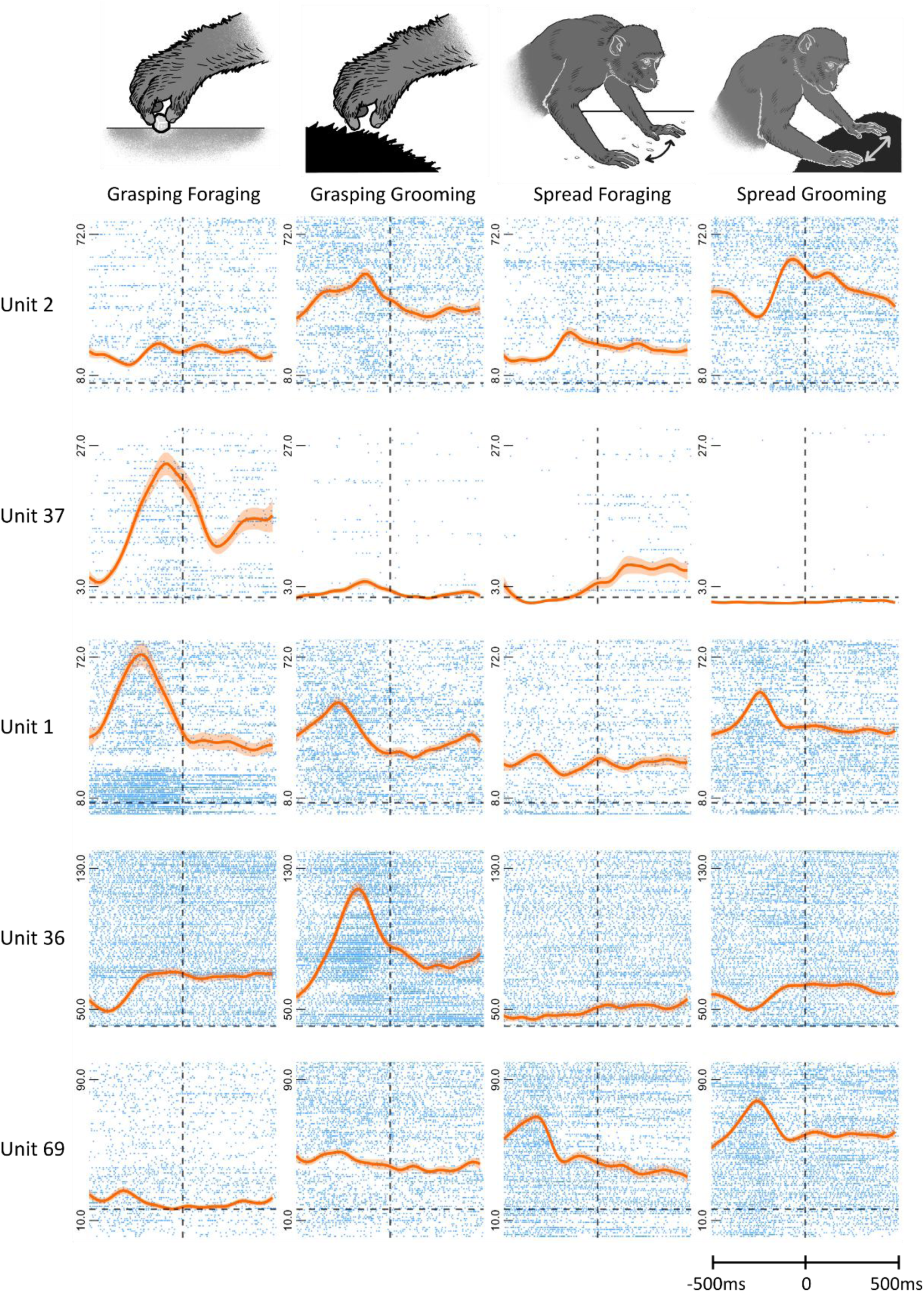
PSTH example units from M2. PSTH and overlaid raster plot of activity of five example units recorded in PMv of M2, showing a variety of response patterns. Mouth activity is not shown because M2 never performed Mouth Grasping.

**Fig. S4.**
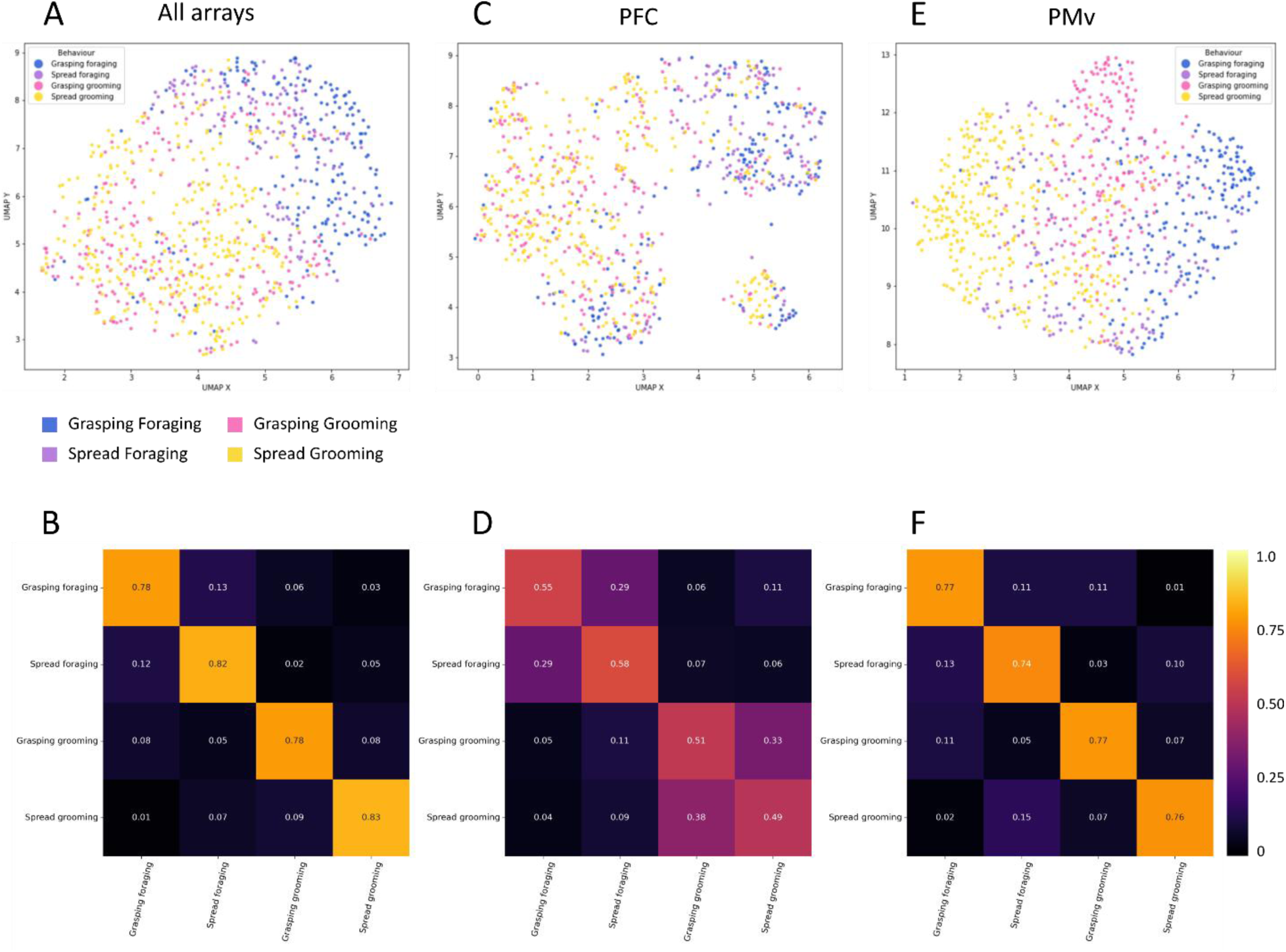
UMAP and XGBoost classifiers for areas combined and separated for M2. (A, C, E) UMAP space embedding of population activity during point-events for M2. (B, D, F) XGBoost classifiers run on the same data. (B) The macro-average F1-score for the classifier run on all data (both PMv and vlPFC areas) is 0.790. (D) For vlPFC only, F1-score = 0.520. (F) For PMv only, F1-score = 0.750.

**Fig. S5.**
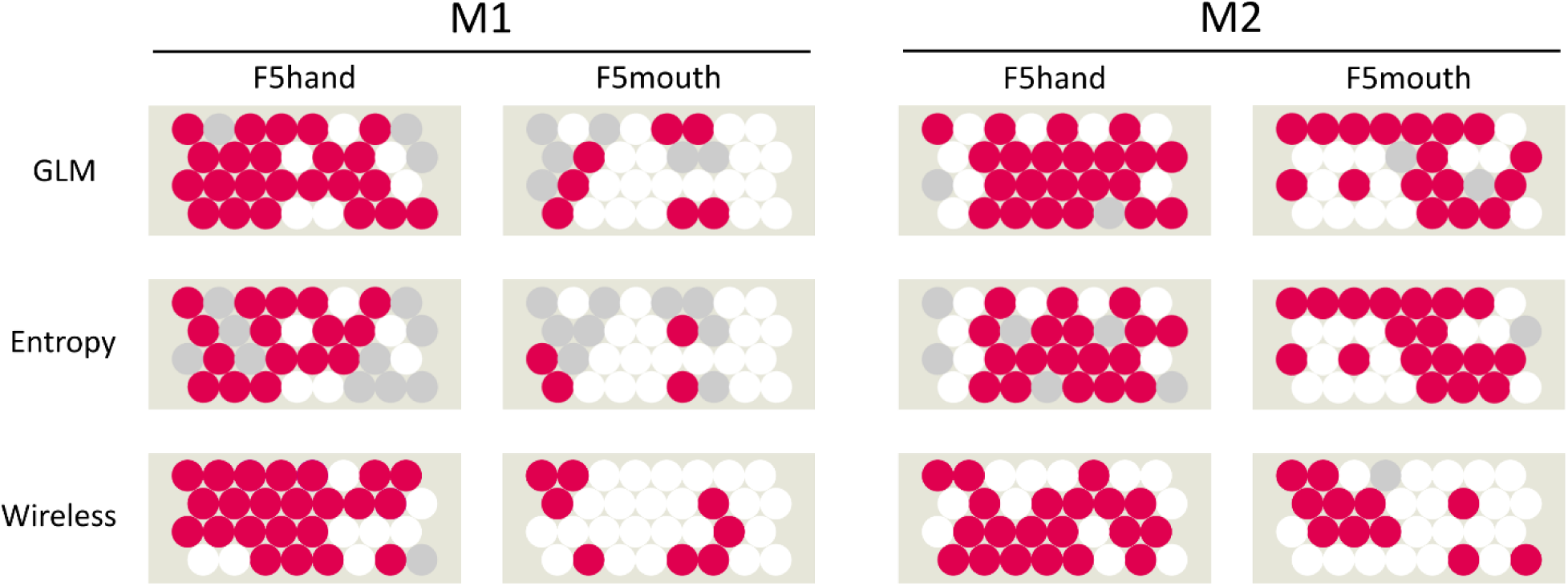
Comparison between GLM and entropy-based test and between tethered and wireless recordings. Electrode maps of FMAs in PMv for M1 (left) and M2 (right). Each circle represents an electrode, colored according to the modulation of units on that channel: red = at least one unit detected on the channel is modulated by grasping, grey = units detected but not showing modulation, white = no units detected on the channel. Top row – modulation is computed with GLM between baseline and grasping epoch in the restrained condition. Bottom row – on the same data, modulation is computed with the entropy-based test. Both statistical procedures are described in Methods. Bottom row – the entropy-based method is applied on grasping (foraging only) events during wireless recording. In the restrained recording, out of a total of 117 units across the two areas and the two monkeys, 66.7% of units were labeled in the same way by the two methods (either both modulated or neither). 25.6% of units were modulated only according to the GLM method, and 7.7% only according to the entropy method. For the comparison between the restrained and wireless conditions, the degree of agreement was computed on the number of channels that showed at least one unit modulated by the behavior. In the head-fixed condition, modulated activity was observed on 30 of 64 channels (46.9%) in M1 and 39 of 64 channels (60.9%) in M2. In the wireless foraging condition, the proportions were highly comparable, with 31 of 64 channels (48.4%) showing modulation in both monkeys.

**Table S1.** Average time budget of each state-based behavior per session. Time budgets (mean across the three sessions ± SD) in seconds and percentages of presence of each state-based behavior across the three sessions in M1 and M2.

| Label | Time budget M1 |  | Time budget M2 |  |
| --- | --- | --- | --- | --- |
|  | (s) | (%) | (s) | (%) |
| Eating | 157.4 ± 69.3 | 2.56 ± 1.13 | 77.1 ± 32.5 | 1.05 ± 0.44 |
| Exploration | 592.1 ± 241.4 | 9.63 ± 3.93 | 99.1 ± 43.0 | 1.35 ± 0.59 |
| Foraging | 1961.7 ± 719.4 | 31.92 ± 11.71 | 158.4 ± 156.3 | 2.16 ± 2.13 |
| Grooming | 193.0 ± 129.7 | 3.14 ± 2.11 | 64.6 | 0.88 |
| LipSmacking | / | / | 132.3 ± 68.4 | 1.80 ± 0.93 |
| Locomotion | 1005.1 ± 555.9 | 16.36 ± 9.05 | 852.7 ± 376.1 | 11.62 ± 5.13 |
| Lying | 38.5 ± 44.2 | 0.63 ± 0.72 | 52.0 ± 51.1 | 0.71 ± 0.70 |
| Mouthing | 27.5 ± 17.6 | 0.45 ± 0.29 | 17.7 ± 25.0 | 0.24 ± 0.34 |
| Presenting | 2.1 ± 3.0 | 0.03 ± 0.05 | 11.3 ± 8.0 | 0.15 ± 0.11 |
| Received Grooming | 2.2 ± 3.1 | 0.04 ± 0.05 | 804.7 ± 172.9 | 10.96 ± 2.36 |
| Resting | 1880.6 ± 371.1 | 30.60 ± 6.04 | 4909.0 ± 325.5 | 66.89 ± 4.44 |
| Scratch | 126.2 ± 39.4 | 2.05 ± 0.64 | 45.4 ± 12.0 | 0.62 ± 0.16 |
| Self-Grooming | 158.9 ± 71.3 | 2.59 ± 1.16 | 40.8 ± 29.7 | 0.56 ± 0.40 |

**Table S2.**
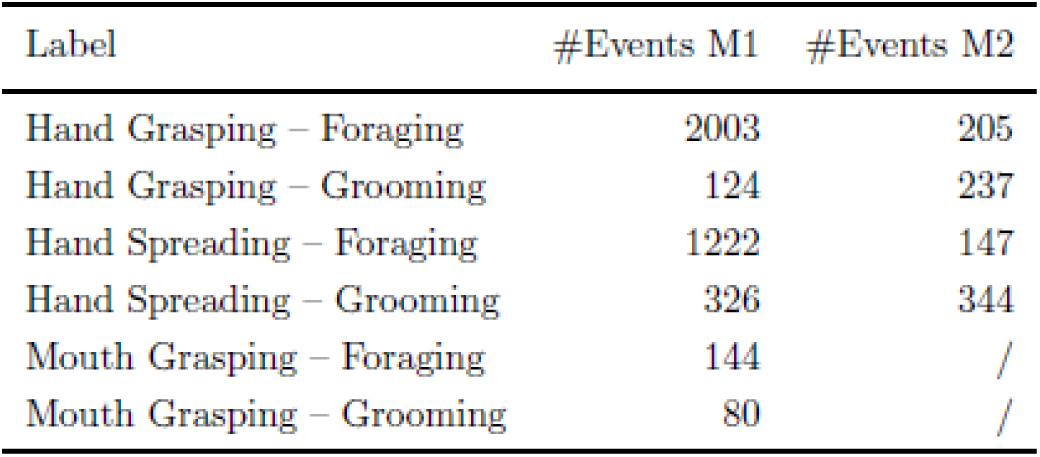
Frequency of all labeled point-based events cumulated across sessions. Frequency of point-based behavioral events involving different motor goals (Grasping, Spreading and Mouth Grasping) and different contexts (Foraging and Grooming) for subjects M1 and M2.

**Table S3.** Numbers of behaviors the units are modulated by. Number of behaviors the units are modulated by, reflecting the degree of mixed selectivity, per each monkey (M1 and M2) and recording area (PMv and vlPFC).

| N Behaviors | M1 |  | M2 |  |
| --- | --- | --- | --- | --- |
|  | PMv | vlPFC | PMv | vlPFC |
| 0 | 1 | 1 | 0 | 6 |
| 1 | 2 | 1 | 2 | 0 |
| 2 | 2 | 1 | 1 | 0 |
| 3 | 15 | 1 | 4 | 3 |
| 4 | 6 | 2 | 30 | 2 |
| 5 | 10 | 3 | 0 | 0 |
| 6 | 6 | 6 | 0 | 0 |

## Notes

### Competing Interest Statement

The authors have declared no competing interest.

